# Mapping the dynamic RNA binding proteome in human effector T cells identifies differentiation and cytotoxicity regulators

**DOI:** 10.64898/2025.12.05.692493

**Authors:** M. Valeria Lattanzio, Koos Rooijers, Kaspar Bresser, Antonia Bradarić, Nandhini Kanagasabesan, Nikolina Šoštarić, Maia Nestor Martin, Nila H. Servaas, Floris P.J. Van Alphen, Arie J. Hoogendijk, Monika C. Wolkers

**Affiliations:** Sanquin Blood Supply Foundation, Department of Research, T cell differentiation lab, Plesmanlaan 125, Amsterdam, The Netherlands, and Landsteiner Laboratory, Amsterdam Institute for Infection & Immunity, Cancer Center Amsterdam-Cancer Immunology, Amsterdam UMC, University of Amsterdam, Meibergdreef 9, Amsterdam; Oncode Institute, Utrecht, The Netherlands; Sanquin Blood Supply Foundation, Department of Research, Bleeding & Hemostasis, Plesmanlaan 125, Amsterdam, The Netherlands

## Abstract

RNA-binding proteins (RBPs) are key regulators of T cell function by controlling (m)RNA fate and fine-tuning protein expression dynamics. Dysregulated RBPs can drive immune diseases and malignancies, highlighting their potential as therapeutic targets. To achieve this, a systematic analysis of the dynamic RBP–RNA interactions is required. Here, we mapped the RNA-binding proteome in human T cells and measured its alterations upon T cell activation using orthogonal organic phase separation (OOPS), analysed with PROMOGEB, a Bayesian linear regression model. This approach uncovered the intricate RNA-binding dynamics of the RBProteome. Gene-editing of such dynamic RNA binders revealed that TUT1 (Star-PAP) maintains the integrity of the T cell differentiation program, and that mutating SF3A1 enhanced the cytotoxic molecule expression and thus target cell killing. Our work provides the most comprehensive analysis of the effector T cell RBProteome to date and shows the potential of identifying RBPs and their binding dynamics as therapeutic agents.

**Teaser:** OOPS analysed with PROMOGEB maps RBP dynamics in human Teff cells, identifying TUT1 and SF3A1 as regulators of T cell fidelity.

## Introduction

T cells are a key component of our defence against infections and malignant cells. Upon antigen encounter, T cells profoundly remodel their gene expression so that they can differentiate, expand, and express key surface receptors and effector molecules, which enable them to kill their target cells. To achieve this, T cells undergo a burst of de novo transcription, together with alterations in translation (1, 2). In addition, translation of pre-existing mRNAs upon activation is initiated (1–3). Combined, these and other features have as consequence that the mRNA and protein abundance in T cells do not fully align, which can be attributed to differential RNA usage, location and translation efficiency, amongst other regulatory features (4, 5). These post-transcriptional regulation (PTR) mechanisms shape the magnitude and temporal coordination of protein expression.

PTR encompasses an intricate network that governs processes like mRNA splicing, stability, decay, localization, and translation, thereby fine-tuning the protein output and ultimately the T cell physiology (6–8). Central to PTR are RNA-binding proteins (RBPs), which recognize target RNAs, mediate RNA modifications, remodel the secondary structure of RNA, and determine the transcript’s fate (9) RBPs are thus indispensable for T cell development, differentiation, and function (6, 10–12). For instance, essential for T cell development are the adenosine-to-inosine editing RBP ADAR1 and the mRNA stability-regulating ZFP36 proteins (13–15). Similarly, the RNA methyltransferase METTL3 regulates T cell lineage commitment and effector function (16, 17). HuR (ELAVL1) boosts cytokine production upon T cell stimulation (2, 11), and Roquin and Regnase1 regulate transcript abundances, including costimulatory receptors, thereby controlling T cell activation and differentiation (12, 18, 19).

Impaired RBP expression or function has been linked to a broad spectrum of autoimmune diseases (14, 20–24). Efforts to develop RBP-direct therapies to boost or curb gene expression are ongoing, for instance by developing small molecules for RBP targeting (25). The first example reaching clinical application is the repair of faulty splicing in spinal muscular atrophy (26, 27). Altering RBP expression to boost T cell responses to infection and tumors has also been tested in preclinical models (10, 11, 28, 29). Yet, despite these important advances, the therapeutic potential of targeting RBPs remains underexplored. A central challenge lies in their mode of action: RBPs are dynamic, multifunctional, and capable of binding diverse RNA substrates. RBPs can also shift their binding according to the cell type and cellular state, as demonstrated in activated T cells (3, 30). Understanding how RBP-RNA interactions are remodeled in a dynamic immune context is therefore paramount for developing effective therapies.

Recent advances in high-throughput technologies, such as crosslinking immunoprecipitation sequencing (CLIP-seq) methods, facilitated the systematic mapping of RBP-target interactions (31). However, these methods focus on individual RBPs and depend on the availability of suitable antibodies. Comprehensive methods that study the global RBP landscape, such as enhanced-RNA interactome capture (eRIC) and orthogonal organic phase separation (OOPS) (32) expanded the RBProteome of Jurkat and CD4+ T cells to up to 1200 RBPs (12, 33). However, because RBP activity is versatile and context-dependent, it is critical to determine how the RNA-binding of RBPs changes upon T cell stimulation, if one wishes to uncover targetable features that modulate T cell functionality.

To address this knowledge gap, we generated a global RBP map with OOPS in resting and stimulated human CD8+ effector T cells (Teff). We developed PROMOGEB, a generalized linear model (GLM) in an empirical Bayes setting, which accounts for our controls as well as missing values, which is inherent to proteomics analysis. We detected 1614 canonical, noncanonical and newly identified RBPs, and we measured their RNA binding activity upon T cell activation. A focused functional CRISPR miniscreen on RBPs displaying altered RNA binding upon T cell activation identified the RNA-editing enzyme PCIF1, the uridylation/polyadenylation factor TUT1 (Star-PAP), and the splicing factor SF3A1 as key regulators of global and gene-specific translation driving T cell differentiation and effector function. Combined, the dynamic RBP map we present here advances our understanding of post-transcriptional networks in adaptive immunity and highlights novel putative targets for therapeutic intervention in T cell-mediated diseases.

## Results

### OOPS optimization in primary human T cells

To comprehensively map the RBP network, we first optimized OOPS for human T cells. OOPS relies on UV-crosslinking of protein-RNA interactions on viable cells and cell lysis with Trizol, followed by acidic guanidinium thiocyanate-phenol-chloroform (AGPC) phase separation to separate RNA-protein complexes from single RNA and protein molecules (**Fig. 1A**). After 3 rounds of AGPC phase separation, RBPs are released from RNA-protein complexes with RNAse and analyzed by immunoblotting and mass spectrometry (MS) analysis. We first defined the best ratio of T cells with Trizol for optimal lysis (10^6^ cells/1ml Trizol; **Fig. S1A**), and the crosslinking efficiency that best preserved cell integrity prior to lysis (30% cross-linking efficiency; **Fig. S1B**). This protocol resulted in effective protein recovery from RNA-protein complexes, as shown for the prototypic RBP HuR (**Fig. S1C, D**).

**Figure 1.**
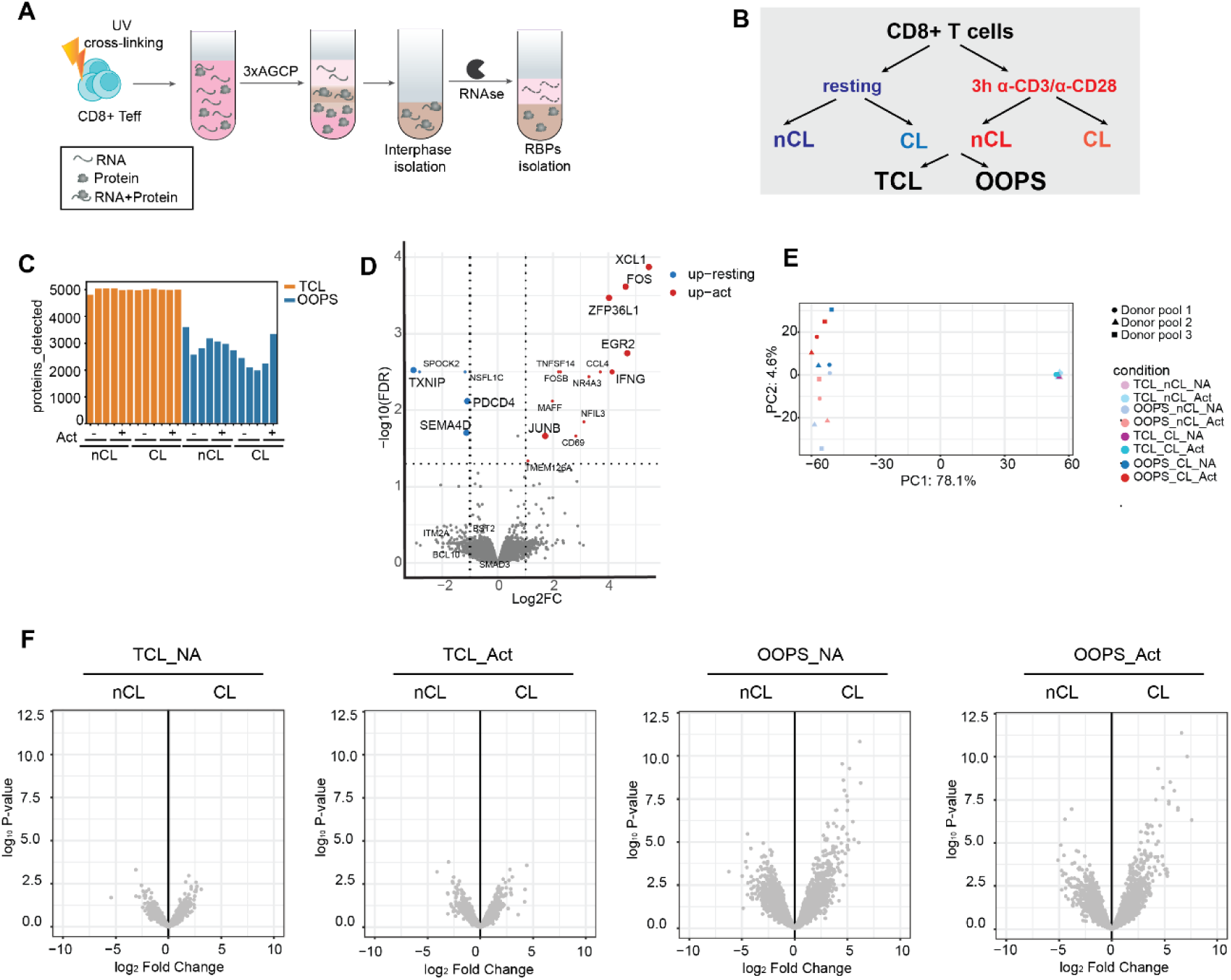
Optimization of OOPS for human T cells. **A** Overview of OOPS. CD8^+^ Teff cells were pelleted and lysed with Trizol. RNA-protein complexes located in the interphase were isolated by 3 rounds of Acid guanidinium thiocyanate-phenol-chloroform extraction (AGCP). RBPs were retrieved by RNase treatment. **B** Experimental scheme depicting generation of Total Cell Lysate (TCL) and OOPS samples, crosslinked (CL) and non-crosslinked (nCL) for each resting and activated (3h with α-CD3/α-CD28) CD8^+^ Teff cells. n=3 donor pools. **C** Number of identified proteins per sample. **D** Differential protein expression (DEP) analysis of TCL samples comparing 3h α-CD3/α-CD28 activation with resting Teff. **E** Principal component analysis of MS data. Treatment conditions are indicated by color, donor pools by shapes. **F** DEP of TCL (left panels) and OOPS samples (right panels), comparing CL with nCL samples.

We next employed OOPS to measure the RBProteome in human T cells. Peripheral-blood derived CD8^+^ T cells were activated for 72h with α-CD3/α-CD28, followed by 7 days of rest to generate Teff cells that rapidly respond to recall responses (*2*). Teff cells were then cultured for 3h in medium (‘resting’; **Fig. 1B**) or were reactivated for 3h with α-CD3/α-CD28, which rapidly induced the production of cytokines (**Fig. S1E)**. To account for background measurements resulting from protein stickiness or carry-over into the interphase, a non-crosslinking (nCL) control was included for each crosslinking (CL) condition (**Fig. 1B**). We prepared total cell lysate (TCL) and OOPS-isolated RNA-protein complex fractions from each sample, resulting in eight experimental conditions (**Fig. 1B**). Using three independent donor pools (≥ 20 donors each), we obtained proteomics data for 24 samples (8 conditions × 3 biological replicates). After applying established quality control workflows and exploratory data analysis (*see Computational Materials and Methods*), around 5000 distinct proteins were identified in the total cell lysates (TCL; **Fig. 1C**). At this early time point of T cell activation, the overall protein expression changes were limited, yet included hallmark early activation-induced proteins such as the transcription factors FOS, JUN, EGR2, and TBX21, the RBP ZFP36L1, and the effector molecules IFN-γ and XCL1/XCL2 (**Fig. 1D)**. Downregulated proteins included, as expected, the chemokine receptor CXCR4, the Thioredoxin-interacting protein TXNIP, and the RBP PDCD4 (**Fig. 1D)**.

We then turned our attention to the RBProteome. Owing to the assay, OOPS samples contained distinctly lower numbers of identified proteins compared to TCL, ranging from 2,000-3,000 identified proteins per OOPS sample (**Fig. 1C; Fig. S1C, F**). For exploratory analysis, we used DEP [v1.24.0] as proteomics data analysis and plotting pipeline. Principal component analysis on normalized, log-transformed data revealed that TCL and OOPS samples appeared in distinct clusters (first principal component; **Fig. 1E**). The second principal component differentiated crosslinked samples (CL) from non-crosslinked samples (nCL) in the OOPS samples (**Fig. 1E**). This was according to our expectation, as retrieval of proteins with RNA affinity is sensitive to crosslinking and thus primarily observed in CL OOPS samples (**Fig. 1F**). This OOPS protocol optimized for T cells thus equipped us to map RBPs in Teff cells and to study their alterations in RNA binding upon T cell activation.

### PROMOGEB, a Bayesian linear regression model to identify RNA-binding proteins

Having optimized OOPS for T cells, we next set up the data analysis pipelines. Our experimental setup was designed to identify up- and down-regulated proteins, RBPs, and changes in RNA-binding (or RNA affinity) upon T cell activation, resulting in 8 distinct experimental conditions (**Fig. 1B**). Established proteomics data analyses pipelines were not suitable for our analyses, because they generally contrast only 2 experimental conditions (*34*). Furthermore, owing to the OOPS methodology, we frequently observed missing values for proteins (**Fig. S1F; Suppl Table 1:** ‘raw data’). Established analysis pipelines calculating logFCs depend on either imputation of missing values, or omission of these missing values. This, however, leads to “infinite logFC values” (when no imputation is used), masking true differences in RNA-binding affinity, or to artificial logFC values (when imputation is used), conflating differences in the proteins’ RNA-binding affinity with expression level differences. Importantly, both methods result in loss of information. Because missing values in proteomics correlate with low abundance (*35*), they inform on the underlying low protein abundance when analyzed in conjunction with other observations. Thus, to include these considerations in the data analysis workflow, we applied generalized linear models (GLMs) in an empirical Bayes setting adapted for proteomics data, which we named PROMOGEB (<u>pro</u>teomics data <u>mo</u>deled with <u>G</u>LM and <u>E</u>mpirical <u>B</u>ayes, **Fig. 2A**). PROMOGEB takes the raw observed intensity values of proteins across all samples, together with a matrix describing the experimental setup (**Fig. 1B**), and outputs linearly independent coefficients dependent on our experimental design, such as OOPS-dependent CL/nCL differences (which we termed RBPness, **Fig. 2B**). As expected, top-ranking proteins in the RBPness coefficient were enriched for the GO term “RNA binding” (**Fig. 2B**). PROMOGEB can thus define the RBPness of proteins in the context of all 8 experimental conditions.

**Figure 2.**
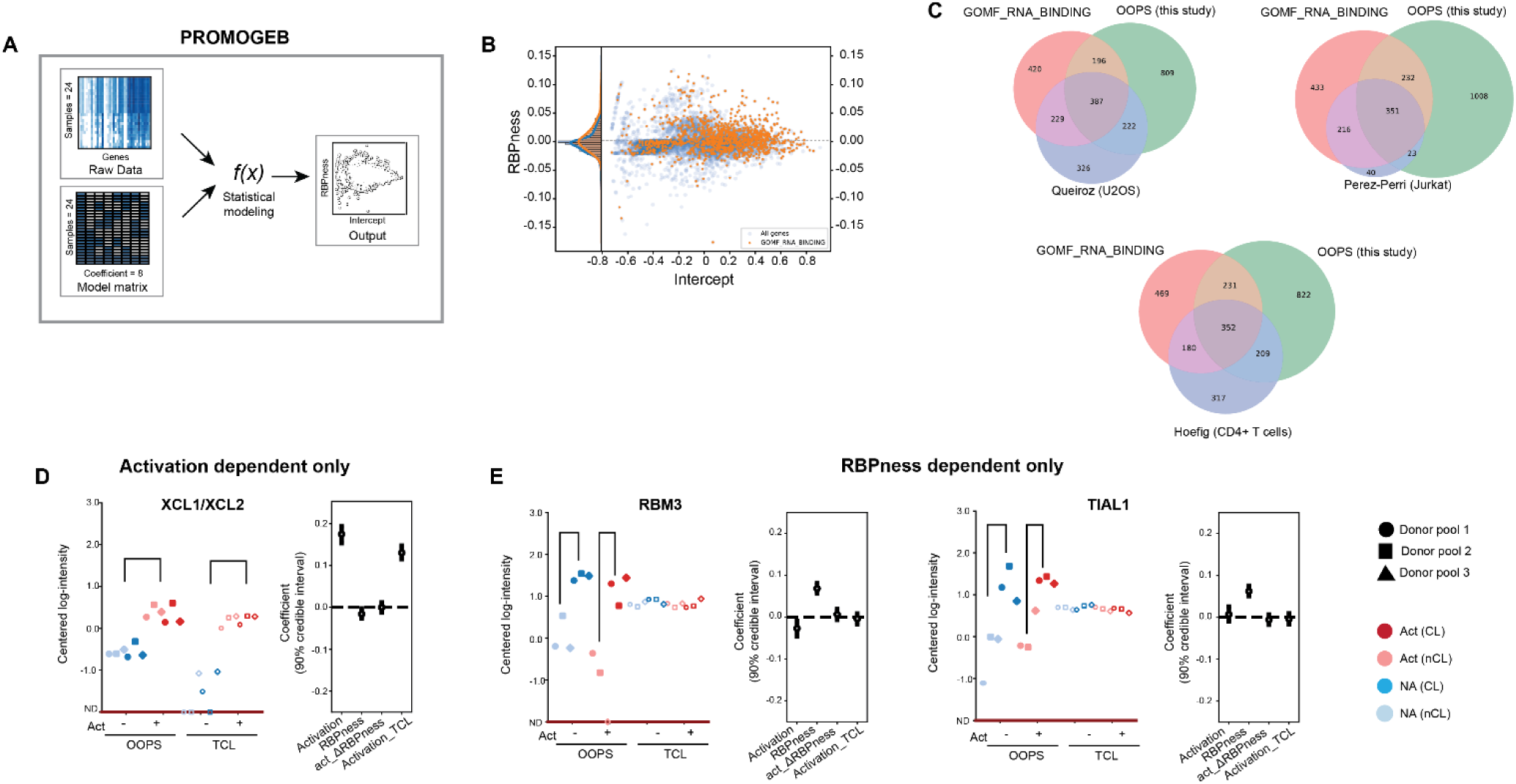
A. Identification of RNA binding proteins by PROMOGEB. **A** Flow diagram of PROMOGEB modeling. Raw proteomics data and design matrix corresponding to experimental setup used as input to our statistical model. PROMOGEB model estimates the most likely (MAP) and range of credible values (HDI) for coefficients. Coefficients express how each gene responds to experimental conditions. **B** MAP estimates intercept (baseline abundance) and RBPness (response in abundance to crosslinking in OOPS samples) coefficients for all proteins (blue) and for proteins annotated with the GO term “molecular function – RNA binding” (orange) that passed our cutoffs. **C** Comparison of RBPs identified in this study (green) with GO term “RNA binding” (red), OOPS from U2O cells (blue; left panel), eRIC from Jurkat cells (blue; right panel), and OOPS from CD4+ T cells (blue, bottom panel). **D-E** Protein abundance indicated as centered-log intensity (left panels) and biological coefficient estimates (HDIs, right panels) for indicated proteins. ND indicates “not detected”. Error bars of coefficients indicate HDI of 90%. Only biologically interpretable coefficients are shown.

### PROMOGEB identifies RBPs with divergent activity and function

To validate the fit of our model, we compared expected and observed missingness rates (**Fig. S2A**). Together with the strong correlation of observed and expected abundances (**Fig. S2B**), PROMOGEB can model protein abundances and their experimental dependencies. We also calculated the logFCs between CL and nCL OOPS samples, using the established DEP workflow, involving imputation of missing values. Comparing the logFCs to our RBPness coefficient showed an overall strong agreement, and outliers were due to missing values (**Fig. S2C)**. PROMOGEB defined 1614 (29% of total detected) proteins as RBPs (*see Computational Materials and Methods*), which substantially overlap and extend published datasets of RBPs in U2OS cells (*32*) and T cell (lines) using OOPS or eRIC (*12*, *33*) (**Fig. 2C**).

Importantly, PROMOGEB separated differential protein abundance along all experimental axes. For instance, the activation coefficient identified the chemokine XCL1/XCL2 in TCL samples (**Fig. 2D**). XCL1/XCL2 detection also increased in CL OOPS samples upon T cell activation, but equally so in the nCL OOPS control samples (**Fig. 2D**). Therefore, XCL1/XCL2 abundance is increased due to T cell activation and is unrelated to RNA binding. Likewise, the prototypic RBPs RBM3 and TIAL1 displayed high abundance in the nCL OOPS control samples (**Fig. 2E**). Yet, these highly abundant proteins are significantly increased in the CL OOPS samples **(Fig. 2E)**, again highlighting the critical contribution of nCL control samples to the data analysis pipeline for well-founded RBP identification (**Fig. S2D)**. Of note, RBM3 and TIAL1 neither varied in protein abundance upon T cell activation, nor did they display altered RNA-binding (**Fig. 2E**), indicating that they belong to the core RBProteome.

We next determined the RBP landscape in Teff cells. Because OOPS is not restricted to poly-A tail-containing mRNAs (32), we identified RBPs annotated as tRNA binders (TRMT1, CTU1, METTL1) and miRNA binders (ZC3HB7, RBM10, TUT7; **Fig. 3A-B**). Consistent with their core function, neither the protein abundance, nor their interaction with RNA changed upon T cell activation (**Fig. 3A-B**). We also found proteins interacting with RNA that were not previously annotated as RBPs. For instance, the protein tyrosine phosphatase PTPRQ was undetectable in TCL but detected as RBP in the CL OOPS samples (**Fig. 3C**). The N-Myristoyltransferase 2 (NMT2) also interacted with RNA independently of T cell activation (**Fig. 3C**). The Thioredoxin interacting protein TXNIP and Utrophin (UTRN) previously annotated as RBPs in cell lines in the RBP database (*32*, *36–38*) were also identified as putative RBPs in Teff cells (**Fig. 3C**). Interestingly, some proteins changed their RBPness upon T cell activation, including HSPA14, which increased its RNA binding without changing its overall protein abundance (**Fig. 3D**). Other RBPs (HAVCR) increased their RBPness or displayed equal RNA binding (SEMA4D), even though their protein abundance was reduced upon T cell activation (**Fig. 3E; Fig. S3A).**

**Figure 3.**
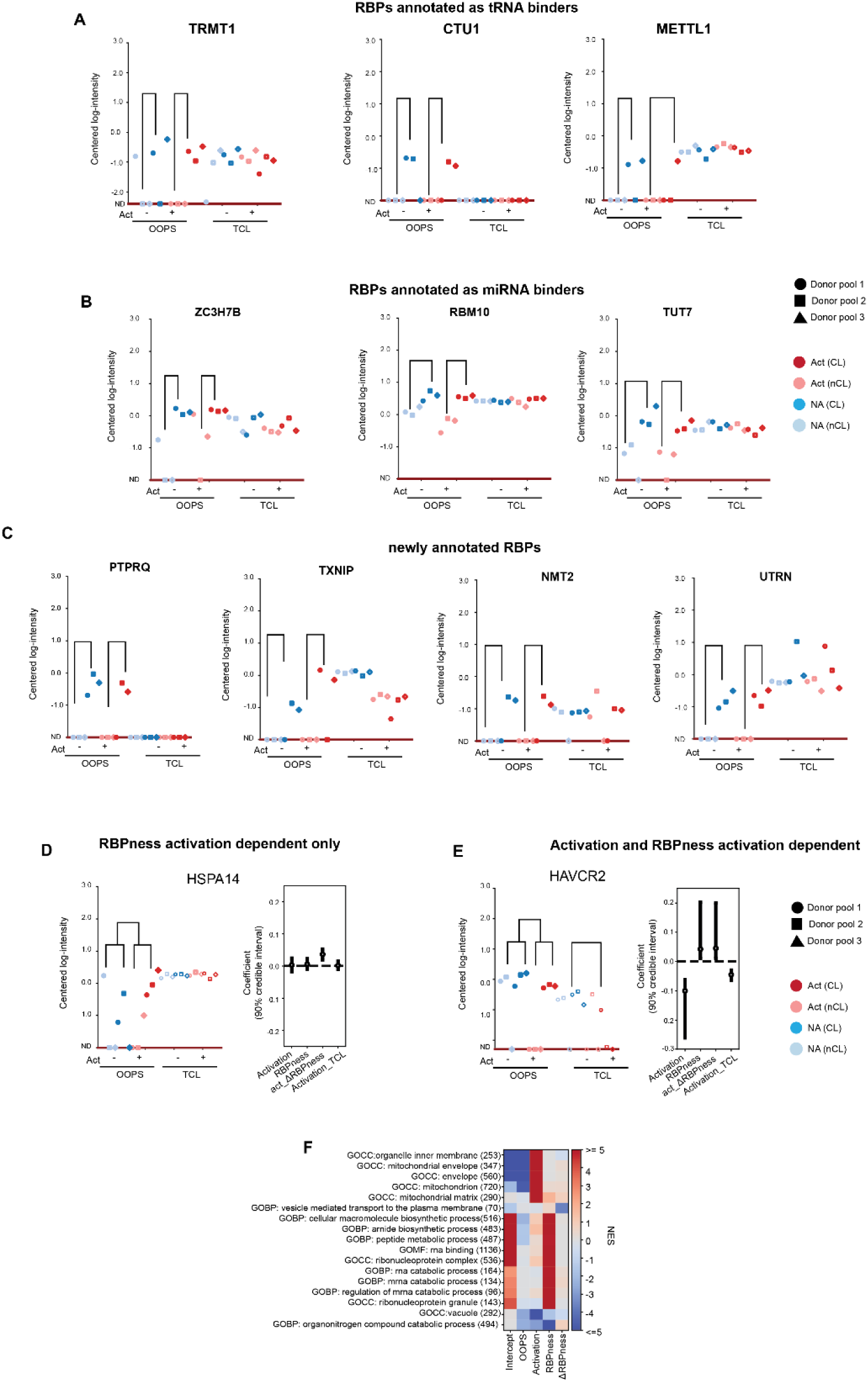
Identification of RBPs interacting with different classes of RNA. **A-C** Proteomics measurements indicated as centered-log intensity for proteins annotated as tRNA (A) and miRNA binders (B), and for proteins newly annotated as RNA binders (C). **D-E** Protein abundance and biological coefficient estimates (HDI) for proteins displaying altered RNA binding upon T cell activation (D) or altered protein expression and RNA binding (E). ND indicates “not detected”. Error bars of coefficients indicate HDI of 90%. **F** Heatmap of normalized enrichment scores (NES) resulting from geneset enrichment analysis (GSEA) for selected components against selected genesets.

Gene set enrichment analysis (GSEA) to all biologically interpretable coefficients that were obtained from PROMOGEB models (**Fig. 3F**) showed, as expected, T cell activation signatures in the activation component and GO terms relating to RNA-binding in the RBPness coefficient (**Fig. S3B**). In conclusion, PROMOGEB obtains credible intervals for the coefficients of interest in the context of 8 experimental settings, most importantly the RNA binding capacity of proteins, and changes thereof upon T cell activation.

### T cell activation drives altered RNA binding of divergent classes of RBPs

RBPs orchestrate all stages of the (m)RNA life cycle, and depending on the signals a cell receives, their binding profile and activity can be modified (*30*, *39*). We therefore asked how RBP-RNA interactions changed upon T cell activation, indicating alterations in the RNA binding activity as ΔRBPness. We zoomed in on RBPs involved in 10 distinct nodules of (post)-transcriptional gene regulation (**Fig. 4**).

**Figure 4.**
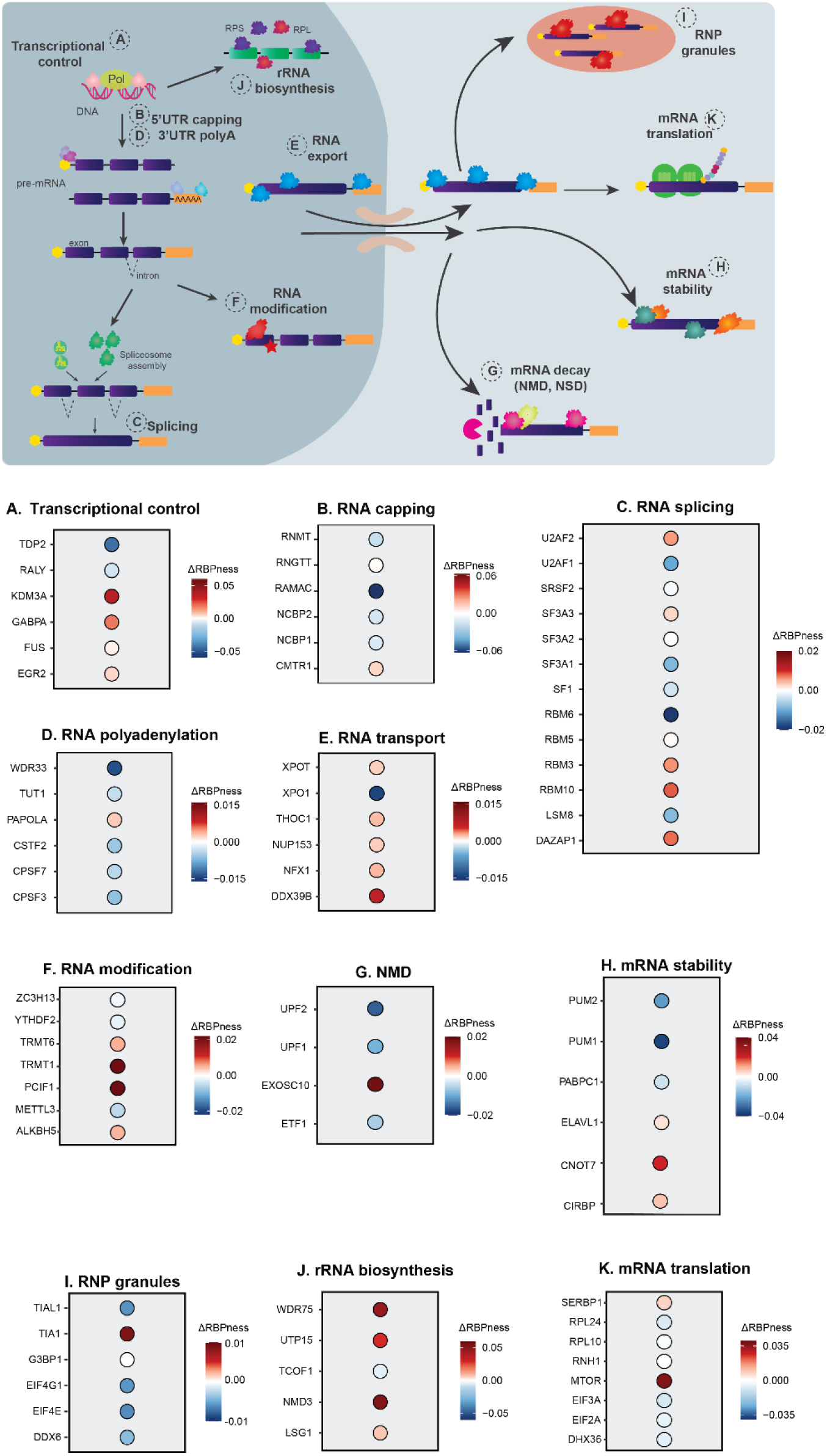
Changes in RBPness upon T cell activation. **A-K** Schematic representation of post-transcriptional mechanisms: (A) transcriptional control, (B) RNA capping, (C) RNA splicing, (D) RNA polyadenylation, (E) RNA transport, (F) RNA modification, (G) NMD, (H) mRNA stability, (I) RNP granules, (J) rRNA biosynthesis, (K) mRNA translation. Each panel depicts the ΔRBPness coefficient, representing the difference in RBPness between resting and activated Teff cells for indicated RBPs.

RBPs involved in transcription control that displayed reduced RNA binding upon T cell activation included the tyrosyl-DNA phosphodiesterase TDP2 and the RNA processing-controlling protein RALY (**Fig. 4A, Suppl Table 1**). Consistent with the transcriptional burst upon T cell activation (*2*), the RNA polymerase controlling protein FUS (*40*) mildly increased its RNA binding (**Fig. 4A**). Notably, the histone demethylase KDM3A and the transcription factors GABPA and EGR2 also interacted with RNA upon T cell activation, suggesting a novel role of these TFs as RNA binders (**Fig. 4A**).

RNA-processing RBPs, including the 5′cap-generating proteins RNGTT, RNMT, and RAMAC and the nuclear cap-binding complex components NCBP1/NCBP2 (*41*) reduced their RNA binding, while the cap-specific m7G methyltransferase CMTR1 (*42*) increased it (**Fig 4B**). This bi-directional change in RNA binding upon T cell activation was also observed for splicing factors: SF1 and U2AF1 decreased, and U2AF2 increased the RNA binding (**Fig 4C**). The SF3A pre-spliceosome complex subunits (*43*) showed a divergent behavior, with decreased (SF3A1), unchanged (SF3A2), or increased (SF3A3) RNA binding, similar to the RBM proteins and DAZAP1 (*44*), RBPs that are involved in alternative splicing (**Fig. 4C**). Core components of polyadenylation (WDR33, CSTF2, CPSF3) and the non-canonical poly(A) polymerase (TUT1) showed decreased RNA binding, but the canonical poly(A)-polymerase PAPOLA increased its RNA binding (*45–48*) (**Fig. 4D**). As to nuclear export, only the export receptor for mature RNAs XPO1 (*49*) strongly reduced its RNA binding, whereas the tRNA transporter XPOT and the core component of the nuclear basket NUP153 (*50*) increased it (**Fig. 4E**). Combined, RBPs involved in RNA processing and nuclear export shift their binding profile to RNA upon T cell activation.

RBPs involved in RNA modifications that displayed reduced RNA binding upon T cell activation included the m⁶A writer METTL3, its regulatory partner ZC3H13, and the m⁶A reader YTHDF2 (**Fig. 4F**). Conversely, increased RNA binding was observed for the m⁶A demethylase ALKBH5, tRNA methyltransferases TRMT6 and TRMT1, and the cap-specific m⁶Am methyltransferase PCIF1 (CAPAM) (*51*, *52*) (**Fig. 4F**). The nonsense-mediated decay (NMD) degradation factors ETF1, UPF1 and UPF2 reduced their RNA binding, just like the poly(A) binding protein PABPC1 and the 3’UTR interactors PUM1/PUM2 (**Fig. 4G, H)**. Yet, the exosome catalytic subunit EXOSC10 and CNOT7 showed increased RBPness (**Fig. 4H),** as did the mRNA-stabilizing and translation regulating proteins ELAVL1 and CIRBP (**Fig. 4H)**. RBPs associated with cytoplasmic granules were unchanged (G3BP1), decreased (TIAL1) or increased (TIA1), and the P-body associated RBP DDX6 (*53*) was decreased, just like EIF4G1 and EIF4E, which recruit mRNA into stress granules to block translation (**Fig. 4I**). Together, RNA binding of RBPs regulates RNA modifications, RNA decay and granule content shifts upon T cell activation.

In concordance with increased translation upon T cell activation (*1*, *54*), core proteins of the pre-rRNA processing machinery WDR75 and UTP15 (*55*, *56*), the ribosome biogenesis-regulating RBPs NMD3 and LSG1 (*57*, *58*) the 40S ribosome regulator RNH1 (*59*, *60*), and the ribosomal proteins RPL24/RPL10 displayed increased RNA binding (**Fig 4J**). This was also true for mTOR, a key factor in T cell development, activation, and translation (*54*, *61*) (**Fig. 4K**). Only the translation initiation factors eIF2A/eIF3A, and the translation-promoting RNA-helicase DHX36 slightly decreased their RNA binding (**Fig. 4K**). In conclusion, T cell activation results in a widespread remodelling of RBP-RNA interactions.

### Identification of RBPs affecting translation in T cells

To study the functional role of RBPs that displayed altered RNA binding upon T cell activation, we performed a mini-screen on 9 candidate RBPs, using CRISPR-CAS9 gene editing (**Fig. 5A, Fig. S4**). The selected RBPs exert different functions, including RNA splicing, modification, stability, transport and translation (**Fig. 4**). RBP deletion in primary T cells was effective (**Fig. S5A**). For SF3A1, we obtained a truncated protein lacking the SURP1 motif, which is essential for its interaction with Splicing factor 1 (SF1) to assemble the canonical pre-spliceosome (*62*) (**Fig. S5A**). All RBP KO/Mut T cells were viable and expanded well throughout cell culture, with only NMD3-KO, SF3A1-Mut, and TUT1-KO T cells slightly lagging behind (**Fig. S5B-C**).

**Figure 5.**
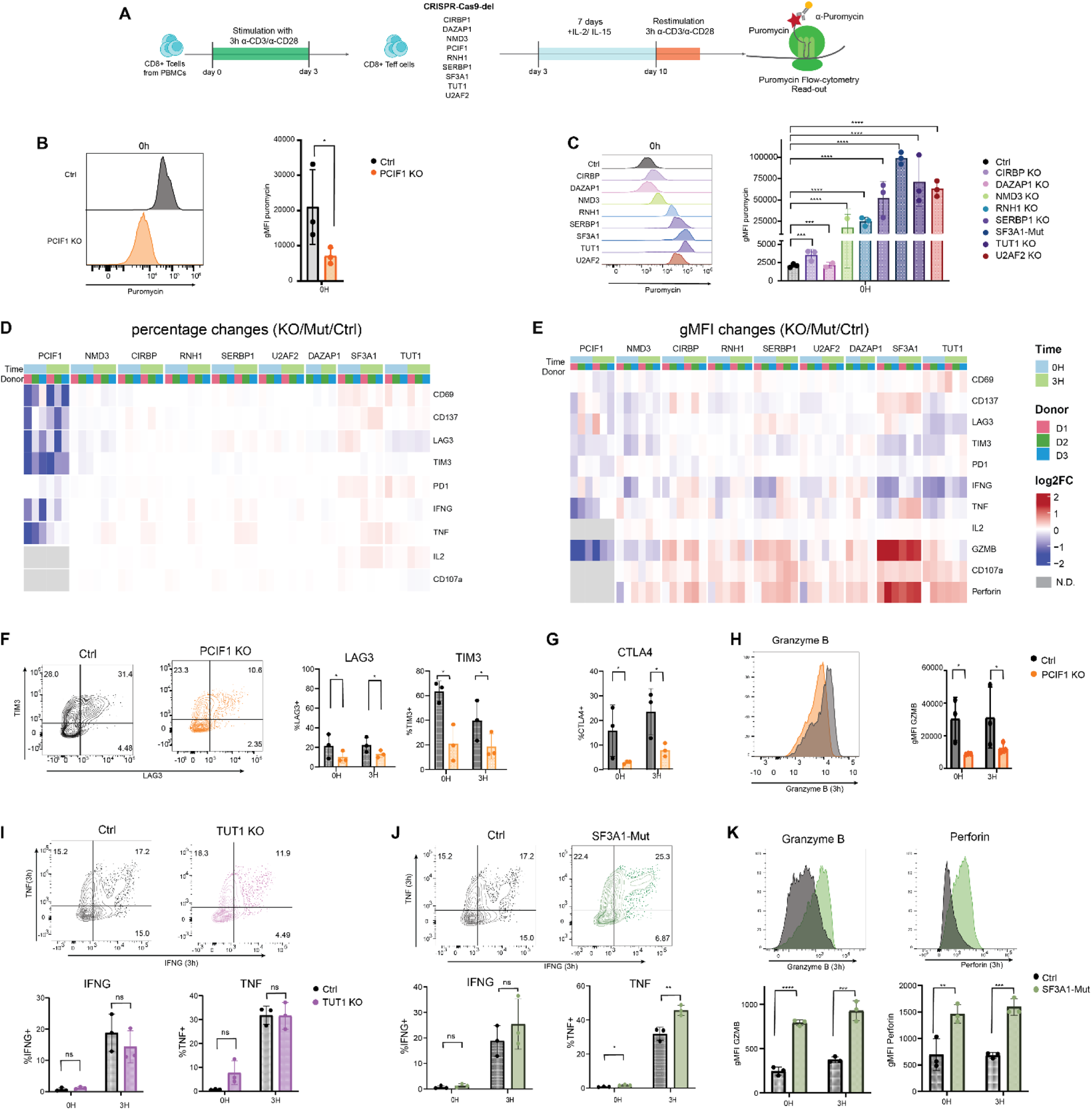
CRISPR-CAS9 gene-editing identifies RBPs modulating T cell function. **A** Schematic overview of the Teff mini-screen. Teff cells generated from PBMCs were nucleofected with Cas9 RNPs targeting the indicated RBPs. After 7 days cell culture, global protein translation was assessed by puromycin incorporation in resting Teff cells. **B.** Representative histogram (left) and gMFI quantification (right) of puromycin expression in PCIF1 KO and Control Teff cells. **C.** Representative histogram (left) and gMFI quantification (right) of puromycin expression in indicated KO Teff cells. **D-E.** Heatmap of log^2^FC of percentage of cells expressing the indicated protein (D), or the gMFI (E). Blue: resting Teff cells, green: Teff cells activated for 3h with α-CD3/α-CD28. N.D. non detected. **F.** Representative LAG-3 and TIM-3 expression in resting PCIF1 KO and control Teff cells (left), and compiled data of the percentage of LAG-3 and TIM-3 expressing Teff cells in resting and 3h activated cells. **G.** Percentage of CTLA4-expressing Teff cells. **H.** Representative GZMB expression (left), and quantification (right) of GZMB gMFI. **I-J.** Representative (top) and compiled data (bottom) of IFN-g and TNF expression in TUT1 KO (I) or SF3A1-Mut (J) and control Teff cells **K.** Representative histogram (top) and compiled data (bottom) of GZMB and Perforin expression in SF3A1-Mut or Control Teff cells. **B-K** Control, CIRBP KO, PCIF1 KO, RNH1 KO, SERBP1 KO, SF3A1-Mut, TUT1 KO, U2AF2 KO data pooled from 3 donors. DAZAP1 and NMD3 KO data pooled from 2 donors. mean ± SD, *p≤0.05, **p≤0.01, ***p≤0.001, ****p≤0.0001 ns: non-significant. Two-tailed ratio unpaired student t-test.

We first measured how RBP deletion affected the global protein translation, using puromycin incorporation as a proxy. 7 out of 9 RBP deletions showed effects. Deleting the 5’Cap m6Am writer PCIF1 substantially reduced puromycin incorporation compared to control T cells, indicating that PCIF1 promotes global translation in T cells (**Fig. 5B**) (*63*, *64*). In contrast, deleting NDM3, RNH1, SERBP1, TUT1, and U2AF2, and mutating SF3A1 increased puromycin incorporation at least a 10-fold, with SF3A1-Mut and TUT1-KO T cells displaying the most pronounced effects (**Fig. 5C**). The changes in global translation also occurred in activated T cells (**Fig. S5D-E**) and did not relate to changes in cell size (**Fig. S5F**), therefore excluding the possibility that differences in puromycin incorporation resulted from morphological changes. Thus, deletion/mutation of 9 RBPs with dynamic interactions detected by OOPS identified one RBP that promotes, and six RBPs that restrain protein synthesis.

### RBP-specific effects on T cell activation-related protein expression

We next studied the effect of RBP deletion on immune-related proteins upon α-CD3/α-CD28 activation. We measured the expression of hallmark T cell activation markers (CD69, CD137, LAG-3, TIM-3, PD-1), cytotoxicity-related molecules (Granzyme B, Perforin, CD107a) and key pro-inflammatory cytokines (IFN-γ, TNF, IL2) by flow cytometry. Deletion of NMD3, CIRBP, DAZAP1, RNH1, U2AF2 and SERBP1 did not show overt effects on percentage (**Fig. 5D**) or expression levels (**Fig. 5E**) of the measured proteins. In contrast, and consistent with the lower overall protein synthesis, PCIF1 deficiency substantially reduced the protein expression, (**Fig. 5D, E**), including that of LAG-3, TIM-3, CTLA-4, and GZMB (**Fig. 5F-H; Fig. S5G**). TUT1-KO T cells had only limited effects on immune-related proteins in this assay, with only Perforin showing slight but significantly increased protein expression (**Fig. 5D-E, 5I; Fig. S5H**). The most prominent effects were observed in SF3A1-Mut T cells (**Fig. 5D, E**). Even though IFN-γ and TNF, as well as PD-1 and LAG3 were only slightly increased (**Fig. 5J; Fig. S5I),** we measured substantially higher GZMB and Perforin expression, together with the surface expression levels of the lysosomal membrane protein CD107a (**Fig. 5K**; **Fig. S5J**), which is indicative of an increased release of cytotoxic molecules. Collectively, these results demonstrate that OOPS can identify RBPs with distinct regulatory activities on T cell effector molecules, with PCIF1 affecting the expression of inhibitory molecules, and SF3A1 regulating cytokine production.

### SF3A1-Mut and TUT1-KO T cells display an altered T cell differentiation profile

The T cell differentiation status is instructed by transcription factors (TF), which define the cell state linked to the production of effector molecules (*65*). To determine if the effects on activation markers and effector molecules observed in Figure 5 resulted from an altered TF expression profile, we measured the expression of key TFs involved in T cell differentiation (TCF1, FOXO1, TOX) and T cell effector function (T-bet, BLIMP1). Notably, even though PCIF1 deficiency impaired the production of effector molecules, TF expression was similar to that of control T cells (**Fig. 6A**), indicating that the diminished production of effector molecules could not be attributed to a different T cell state. In contrast, SF3A1-Mut T cells expressed higher levels of TOX (**Fig. 6B**), a TF that regulates T cell persistence. SF3A1 mutation also resulted in increased T-bet expression, the master regulator of effector differentiation (**Fig. 6C**) (*65*), which corresponds with the observed increased production of effector molecules (**Fig. 5K; Fig. S5J**). T-bet directly antagonizes the expression of TCF1, the T cell differentiation and memory fate factor (*66*). However, TCF1 expression remained unchanged in SF3A1-Mut T cells (**Fig. S6A)**. In addition, CD27 and CCR7 expression was substantially downregulated in SF3A1-Mut T cells, which is indicative of terminal effector differentiation (*67*) (**Fig. 6D**), which points to diminished memory-like features in SF3A1-Mut T cells.

**Figure 6.**
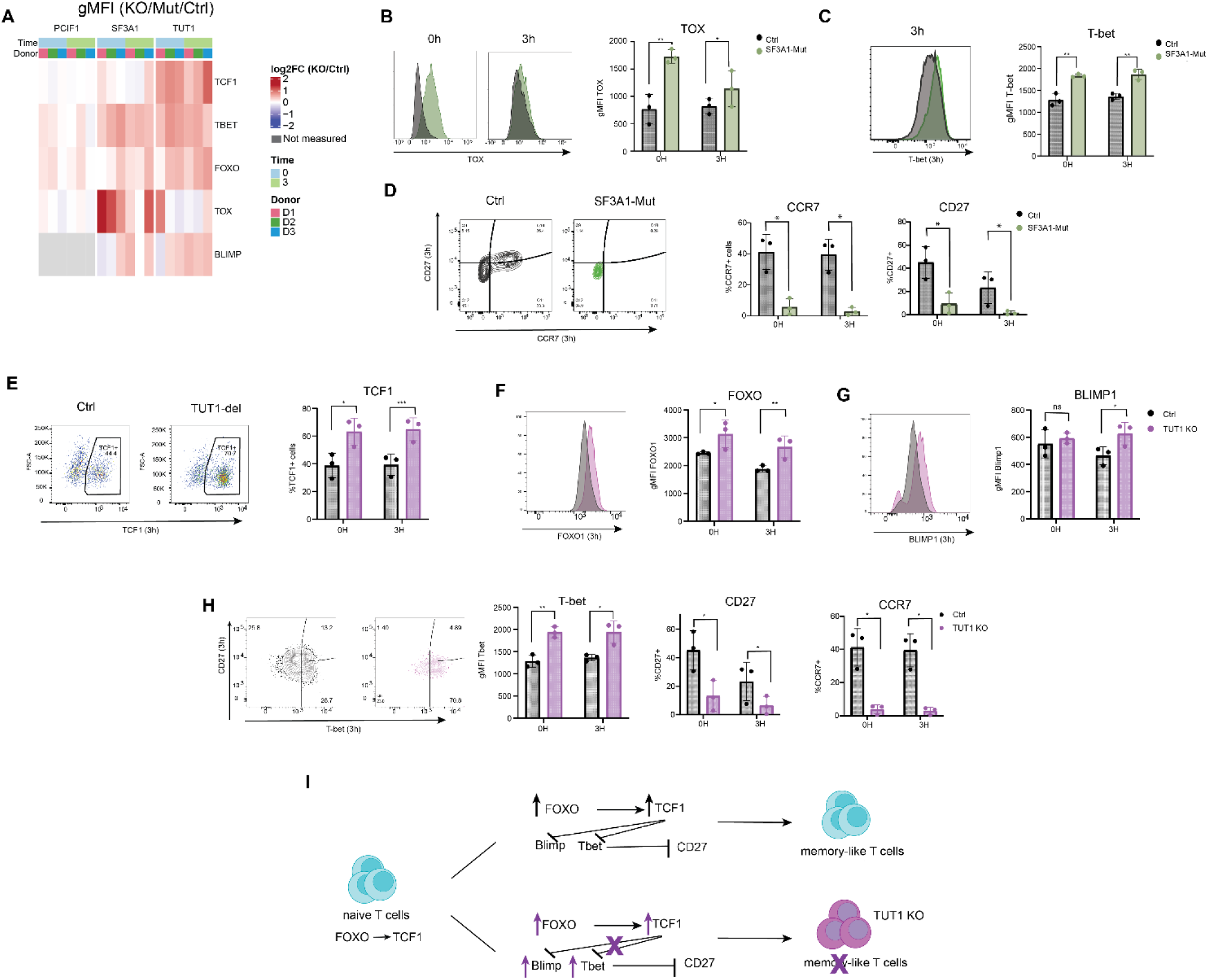
TUT1 deletion alters the T cell differentiation profile. **A.** Heatmap depicting the log_2_FC gMFI expression of transcription factor expression between Control and indicated KO/Mut Teff cells. **B-D.** TOX expression (B), T-bet expression (C) and CCR7 and CD27 expression (D) in SF3A1-Mut and Control Teff cells. **E-H** TCF1 expression (E), FOXO1 expression (F), BLIMP1expression (G), and T-bet, CD27 and CCR7 expression (H) in TUT1 KO and Control Teff cells. **I.** Scheme of effects on T cell differentiation-associated proteins in TUT1 KO Teff cells. **B-H** Data pooled from 3 donors. mean ± SD, p≤0.05, **p≤0.01, ***p≤0.001, ****p≤0.0001 ns: non-significant. Two-tailed ratio unpaired student t-test.

We then turned our attention to TUT1 KO T cells. Intriguingly, TUT1 deletion resulted in higher TCF1 expression levels in both resting and activated T cells (**Fig. 6E; Fig. S6B**). In agreement, the expression of FOXO1, which directly promotes TCF1 expression (*68*), was also increased in TUT1 KO T cells (**Fig. 6F**). Elevated FOXO1 and TCF1 expression can bias T cell differentiation towards a memory-like phenotype, resulting in reduced expression of the effector-driving transcription factors T-bet and BLIMP1 (*65*). Unexpectedly, however, T-bet and BLIMP1 expression was increased in TUT1 KO cells (**Fig. 6G, H**), concomitant with a substantial loss of the memory state-associated proteins CD27 and CCR7 (**Fig. 6H; Fig. S6C**). To determine whether the uncoupling of TCF1 expression from T-bet expression depended on the timepoint of gene-editing, we deleted TUT1 a week later in Teff cells, i.e. on day 10 of cell culture (**Fig. S6D**). The increased TCF1 expression in TUT1 KO T cells was also maintained at that time point (**Fig. S6E**). Yet, the effects on FOXO1, Tbet, and CD27 were by and large lost (**Fig. S6F-G**), indicating that the effect of TUT1 deletion on T cell differentiation is time-dependent, except for TCF1. Because TCF1 was always affected, we asked whether TUT1 directly regulated TCF1 expression through its polyadenylation activity. The overall *TCF7* mRNA levels (encoding TCF1) did not change in TUT1 KO T cells (**Fig. S6H**). Yet, in two out of three tested donors, the polyadenylation pattern of *TCF7* altered in TUT1 KO T cells, but not that of *GAPDH* (**Fig. S6H)**, suggesting that *TCF7* mRNA is a direct target of TUT1. Combined, these data show that TUT1 orchestrates the TF expression during T cell differentiation (**Fig. 6I**).

### SF3A1 deletion boosts T cell cytotoxicity

Lastly, we investigated the activity of PCIF1 KO, TUT1 KO, SF3A1-Mut T cells when exposed to target cells. We co-cultured gene-edited T cells with the melanoma cell line Mel888, which we engineered to express an α-CD3 single-chain fragment variable antibody at the cell surface, enabling us to study T cell responses in the context of cell-cell interactions (**Fig. 7A**). We first measured the expression of activation markers and effector molecules upon 16h exposure to the target cells. PCIF-KO T cells showed no or very modest effects (**Fig. 7B-F; Fig. S7A, B**). In contrast, whereas SF3A1-Mut and TUT1 KO T cells expressed only slightly higher levels of the activation markers CD25, CD69 and CD137, the cytokine expression was markedly increased (**Fig. 7B-D; Fig. S7A, B**). SF3A1-Mut and TUT1 KO T cells also expressed higher levels of GZMB and Perforin, with SF3A1-Mut T cells displaying the highest production levels of these cytotoxic molecules (**Fig. 7E, F**).

**Figure 7.**
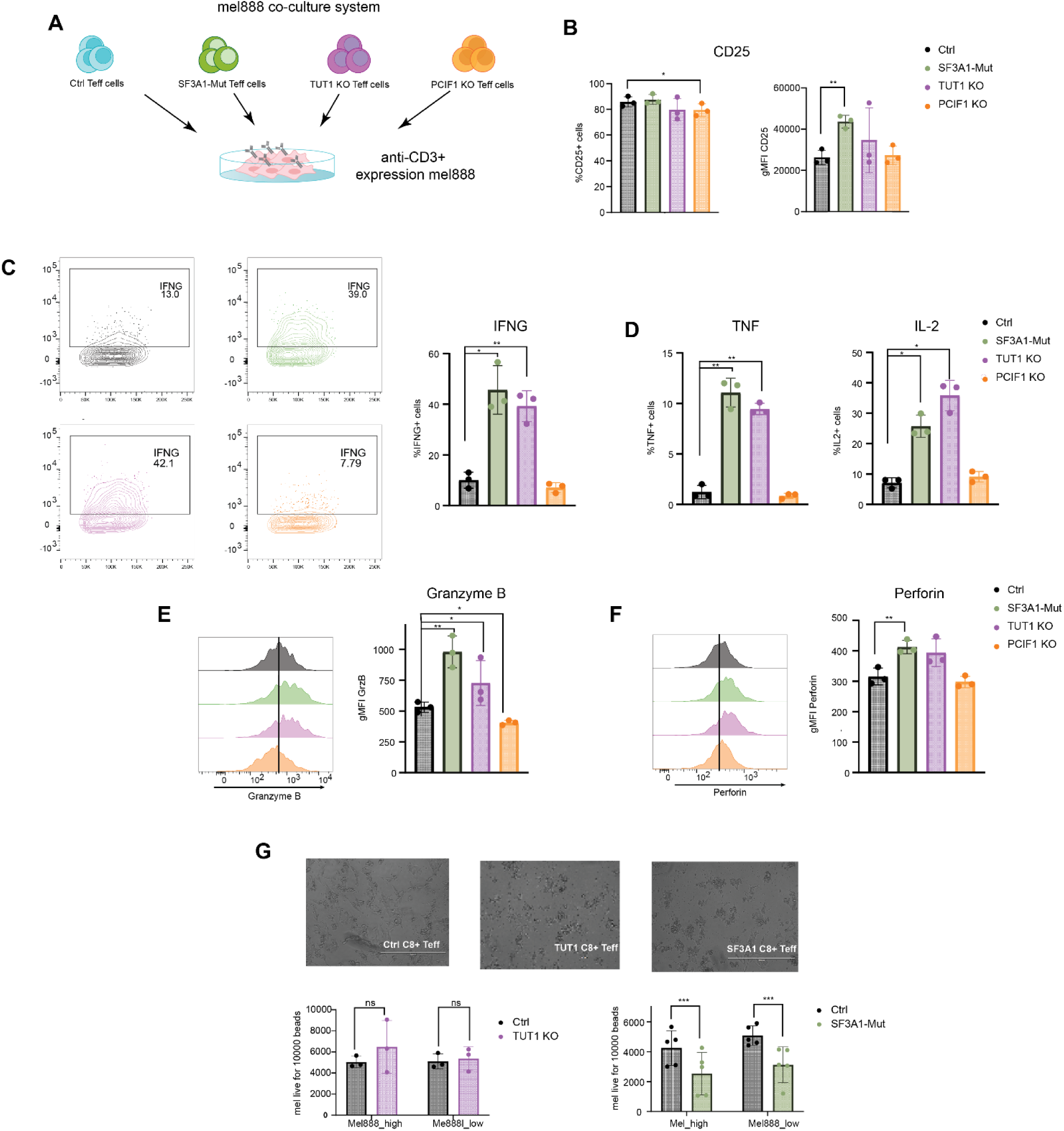
SF3A1 mutation boosts T cell cytotoxicity. **A.** Scheme of T cell co-culture of indicated KO/Mut Teff cells with α-CD3 (high and low level)-expressing Mel888 tumor cells. **B.** Percentage and gMFI of CD25 expression upon 16h of co-culture with α-CD3^+^ medium Mel888 cells. **C-D.** Percentage of IFN-g (C), TNF, and IL2 (E) -producing Teff cells. **E-F.** gMFI quantification of GZMB (E) and Perforin (F) expression. **G.** Representative image after 16hrs of co-culture of Control, TUT1 KO and SF3A1-Mut Teff cells with α-CD3^+^ high Mel888 cells (top, scale bar = 400μm). Quantification (bottom) of live Mel888 cells per 10.000 counting beads in Control, TUT1 KO and SF3A1-Mut Teff cells co-culture. **B-G** Data pooled from 3 donors (**B-F**). Control and TUT1 KO data pooled from 3 donors (**G**). Control and SF3A1-Mut data pooled from 5 donors (**G**). mean ± SD, p≤0.05, **p≤0.01, ***p≤0.001, ****p≤0.0001 ns: non-significant. Two-tailed ratio unpaired student t-test.

We next questioned how RBP deletion modulated the capacity of Teff cells to kill target cells. As expected from its limited effects on effector molecule expression, PCIF1 KO T cells did not improve tumor cell killing compared to control T cells (**Fig. S7C**). Likewise, and irrespective of their higher cytokine expression and increased expression of cytotoxic molecules (**Fig. 7C-F; Fig. S7B**), also TUT1KO T cells did not kill Mel888 cells better, whether Mel-888 cells expressed high or low levels of α-CD3 (**Fig. 7G**). In sharp contrast, SF3A1-Mut Teff cells more potently killed Mel888 cells than control T cells (**Fig. 7G**). In sharp contrast, and in alignment with SF3A1-Mut Teff expressing the highest levels of activation markers and effector molecules, we found that SF3A1-Mut Teff cells more potently killed Mel888 cells than control T cells. In conclusion, we show here how measuring the dynamic RBProteome helps uncover novel regulatory nodes in T cells.

## Discussion

In this study, we present a comprehensive map of the RBProteome in human Teff cells and show its versatility upon T cell activation. By optimizing OOPS for T cells and by generating a new analysis pipeline, PROMOGEB, we identified 1614 known and new RBPs, which expand the currently available RBP maps in T cell lines (*12*, *33*).

A key advantage of using PROMOGEB for data analysis is that it directly extracts the RNA-binding activity derived from OOPS. By using background measurements available through our experimental setup (e.g. using non-crosslinked and TCL samples), PROMOGEB could delineate RNA-binding, protein-intrinsic chemical properties, and changes upon T cell activation. Using this approach, we identified canonical and non-canonical RBPs such as PTPRQ and TXNIP, but also transcription factors such as EGR2 and chromatin regulators. Even though their moonlighting RBP activity awaits experimental confirmation, our findings expand the current repertoire of dual - putative and confirmed-DNA-RNA binding proteins (*69*, *70*).

OOPS recovers RBP binding not only from poly-A containing mRNA but from all RNA classes (32). Therefore, we also detected tRNA- and miRNA-binders. Furthermore, the observed alterations on RBPness upon T cell activation may help identify how target specificity of RBPs is governed. For instance, the pre-mRNA splicing factor U2AF2 gains target selectivity through interaction with its partner U2AF1, which facilitates U2AF2 binding to weaker pyrimidine-tracts at the 3′ splice site (*71*). With U2AF1 showing reduced RNA binding upon T cell activation, it is therefore tempting to speculate that the target selectivity of U2AF2 changes, together with its observed higher RNA binding.

Follow-up studies on RBPs with altered RBP binding in activated T cells identified SF3A1 as a putative therapeutic target for boosting T cell responses. How SF3A1 regulates T cell cytotoxicity is yet to be determined. SF3A1 is a core subunit of the SF3a complex (*62*). The SURP1 domain deleted in SF3A1-Mut T cells interacts with splicing factor 1 (SF1), which is required for the assembly of the pre-mRNA splicing complex (62, 72). Intriguingly, SFSWAP and CHERP, two other SURP1-containing proteins involved in alternative splicing can also engage with SF1 (62). It is therefore conceivable that disabling the interaction of SF3A1 with SF1 promotes alternative splicing. Notably, alternative splicing is linked to T cell activation and differentiation (73). Indeed, some effector molecules that were increased in SF3A1-Mut T cells were previously shown to be regulated through splicing, such as TNF (74). However, provided the breadth of observed effects, we postulate that SF3A1 mutation rather acts upstream of effector molecules, e.g. by boosting the expression of T-bet. Notably, SF3A1-Mut T cells also showed substantial increases in global translation. Whether this is due to indirect effects, for example via changes in NMD, mRNA export, or the expression of translation regulators, is yet to be defined. In conclusion, our results position SF3A1 as a gatekeeper of T cell effector function and highlight its potential as a therapeutic target to enhance anti-tumor activity.

Surprisingly, even though TUT1 KO T cells produced similar levels of effector molecules as SF3A1-Mut cells when exposed to target cells, TUT1 deletion failed to boost target cell killing. TUT1 deletion disrupted the T cell differentiation program, as evidenced by increasing not only TCF1 and FOXO1 expression, promoting a memory-like state (75, 76), but also T-bet and BLIMP1, markers of terminal differentiation (76). This decoupling of differentiation programs was observed only when TUT1 was deleted early on, but not at later time points, suggesting that TUT1 is essential to guide early lineage decisions. Mechanistically, our data indicate that TUT1 can regulate *TCF7* mRNA polyadenylation, which extends this concept of post-transcriptional control through polyadenylation to key transcription factors.

Importantly, the global analysis on RBPness we present here highlights the intricate alterations in RNA-protein interactions upon T cell activation. Perhaps not surprisingly, alterations in RBPness are rather observed for specific proteins than for a protein group belonging to specific biological processes. For instance, the bi-directional changes in RNA binding of RNA methylation writers and readers may reflect a shift in the methylation profiles upon T cell activation, as recently reported (*17*). Likewise, PCIF1 increases its RBPness upon T cell activation, and PCIF1 depletion uncovered its key role in promoting global translation. Whereas our study confirms the reported link of PCIF1 with translation regulation (*51*, *52*, *77*) other studies found that PCIF1 regulates mRNA stability (78), yet which depended on the organ analysed in mice (*64*). These findings highlight the context-specific activity of PCIF1. Collectively, in human Teff cells, PCIF1 promotes translation, exemplifying how RNA modifications contribute to T cell function. In conclusion, our study establishes OOPS and PROMOGEB analysis as powerful tools to determine the dynamic RBProteome in T cells, and it showcases how it can help identify candidates with potential therapeutic relevance. Applying this to other immune cell types should advance our understanding of the context-dependent functions of RBPs.

## Materials and Methods

### T cell activation and cell culture

Human T cells from anonymized healthy donors were used after written informed consent (Sanquin), in accordance with the Declaration of Helsinki (Seventh Revision, 2013). Peripheral blood mononuclear cells (PBMCs) were isolated by Lymphoprep density gradient separation (Stemcell Technologies). To generate effector T cells (Teff), CD8^+^ T cells were enriched from cryopreserved, defrosted PBMCs with the CD8^+^ T cell isolation kit (Miltenyi Biotec, purity >85%). T cells were activated for 72 h with 1 μg/mL plate-bound α-CD3 (HIT3a, Biolegend) and 1 μg/mL soluble α-CD28 (CD28.2; Biolegend), as previously described (*2*). Cells were cultured at 37°C, 5% CO2 in culture medium (IMDM, GIBCO, Thermo Fisher Scientific, supplemented with 10% fetal bovine serum (FBS), 100 U/mL penicillin, 100 μg/mL streptomycin, 2 mM L-glutamine). After activation, Teff cells were harvested and cultured at a density of 1,5×10^6^/mL for 7 days in standing T25/75 tissue culture flasks (Thermo Scientific) in the presence of 100 IU/mL recombinant human (rh) IL-2 (Proleukin). Culture medium was refreshed every 2 days. Upon nucleofection, cells were cultured for 3d in T cell mixed media (Miltenyi Biotec) supplemented with 5% heat-inactivated human serum, 5% FBS, 100 U/mL Penicillin, 100 μg/mL streptomycin, 2 mM L-glutamine, 100 IU/mL rhIL-2, before switched back to culture medium with 100 IU/mL rhIL-2.

### Orthogonal Organic Phase Separation

OOPS methodology (*32*) was adjusted to Teff cells as follows: 50×10^6^ cells/sample were cultured in a T25 flask with 1ug/ml α-CD3 (Pelicluster CD3, Sanquin) and 1 μg/mL α-CD28 in culture media for 3h. Non-activated Teff cells served as control. Cells were washed twice with ice-cold PBS. For crosslinking (CL), cells were resuspended in 4ml ice-cold PBS, plated on a 150mm Petri-dish (Nunc™ Petri Dishes, Thermo Scientific) and cross-linked at 254nm with 150 mJ/cm^2^ using the Stratalinker UV-Crosslinker (during protocol optimization: 2 rounds of 400 mJ/cm^2^ followed by one round of 200 mJ/cm^2^). Cells were collected in 5ml ice-cold PBS by careful scraping, washed and pelleted. Dried pellets were homogenized in 5ml acidic guanidinium-thiocyanate-phenol (Trizol, Thermo Fisher Scientific) in protein-low binding tubes (Eppendorf) and stored at -80°C. To extract RNA-protein-complexes, 1ml chloroform (1:5 ratio Chloroform:Trizol) was added upon thawing, vortexed at maximum speed and centrifuged for 15min at 12,000g at 4°C. Aqueous and organic phases were carefully discarded. RNA-protein complex extraction from the interphase was repeated twice for further purification. Protein precipitation from the interphase was performed by adding 9:1 (vol/vol methanol:sample) methanol, vortexed at maximum speed, centrifuged for 10min at 14,000g at 4°C. Supernatant was removed and pellet stored at -20°C for 24hrs. RNA-binding protein recovery was performed by resuspending pellets in 100mM TEAB buffer (Sigma-Aldrich) and 1% of SDS (Sigma-Aldrich), and sonicating in ice-cold water for 15min (Branson 2200 Ultrasonic Cleaner), before heating samples for 20min at 95°C and cooling on ice for 2min. Samples were treated with 4ug RNase A/ 1U RNase T1 (both Thermo Scientific) for 4h at 37°C, homogenized in 1ml Trizol and 200ul Chloroform and centrifuged for 15min at 12,000g at 4°C. The organic phase containing proteins was collected and transferred to a new protein low-binding tube. Proteins were precipitated by adding 1350μl ice-cold methanol per 150μl organic phase and centrifuged at maximum speed for 10min at 4°C. Precipitated proteins were washed with 1ml ice-cold methanol, and pellets were stored at -20°C for up to 1 month.

### Sample preparation for MS

Peptide samples were desalted using in house prepared Empore C18 STAGE tips. Peptides were eluted with 0.5% (v/v) acetic acid, 80% (v/v) acetonitrile. Sample volume was reduced by SpeedVac and supplemented with 2% acetonitrile, 0.1% TFA to a final volume of 4.5 μl. 3 μl of each sample was injected for MS analysis on a Orbitrap Fusion Lumos, coupled to an Ultimate 3000 Nano HPLC system (Thermo Scientific). Eluted peptides were separated using in house prepared columns (New Objective type FS360-75-8-N-5-C20, Inc., Woburn, MA, USA) filled with 1.9 μm C18 particles (Dr.Maisch, Ammerbuch-Entringen, Germany) at a flow rate of 300 nL/min, Buffer A was composed of 0.1% formic acid, buffer B of 0.1% formic acid, 80% acetonitrile (BioSolve, France). Peptides were loaded for 17 min at 300 nl/min at 5% buffer B, equilibrated for 5 min at 5% buffer B (17-22 min) and eluted by increasing buffer B from 5-15% (22-87 min) and 15-38% (87-147 min), followed by a 10 min wash to 90% and a 5 min regeneration to 5%.

### LC-MS/MS

Mass spectrometry (MS) was run in DDA mode. Survey scans of peptide precursors from 400 to 1500 *m*/*z* were performed at 120K resolution (at 200 *m*/*z*) with a 1.5 × 10^5^ ion count target. Tandem MS was performed by isolation with the quadrupole with isolation window 1.6, HCD fragmentation with normalized collision energy of 30, and rapid scan MS analysis in the ion trap. The MS^2^ ion count target was set to 1.5 x 10^4^ and the max injection time was 35 ms. Only precursors with charge state 2–7 were sampled for MS^2^. The dynamic exclusion duration was set to 60s with a 10ppm tolerance around selected precursor and its isotopes. Monoisotopic precursor selection was turned on. The instrument was run in top speed mode with 3s cycles. All data were acquired with SII for Xcalibur software.

### Parsimonious gene identification from peptides and sample integration

To integrate observations across all batches of protein samples, we integrated observed peptides across all measured samples. We then used graph algorithms (min-cost-max-flow) to identify the parsimonious set of UniProt proteins explaining the observed peptides. The min-cost-max-flow problem was constrained such that annotated UniProt entries (SwissProt) were preferred over computationally annotated entries (TrEMBL). Also, preference was given to proteins with high evidence level (UniProt field ‘proteinexistenc’). Implementation can be found at: https://github.com/krooijers-sanquin/2024-proteomics-preprocessing. Clusters of UniProt entries explanatory for a set of peptides were further reduced to their associated gene (HGNC ID); in most cases leading to a single unique gene identifier. We then referred back to observed data in the proteomics samples and summed observed intensities for each peptide across all peptides for each gene identifier. We then obtained a matrix of genes per sample, which was used for downstream analyses. UniProt data were downloaded as XML file from https://ftp.uniprot.org/pub/databases/uniprot/current_release/knowledgebase/reference_proteomes/Eukaryota/UP000005640/ at 4^th^ of October, 2023 (207933 entries).

### Linear modeling and Bayesian regression on proteomics data, PROMOGEB

Our experimental setup exists of a fully crossed-design matrix across the factors “library type” (either OOPS or TCL), “crosslinked” (either “yes” or “no”) and “activated” (either “yes” or “no”). We employed an empirical Bayesian modeling approach, where output intensities are modeled as originating from a hurdle-log-normal distribution. This distribution has 3 parameters: mu (expected value), tau (scale parameter) and psi (probability of observing the value, versus it being missing). These parameters are further modeled in a hierarchical fashion as outcomes of biological and technical parameters. The biological parameters link the experimental setup via a design matrix to a coefficient matrix, where each gene is attributed a set of coefficients. The technical parameters relate abundance to probability of missing values (unique for each sample) and scaling of measured intensity values (unique for each sample). We obtained the technical parameters for all samples using a single fit of likelihood distribution by identifying the maximum a posteriori (MAP) via an adam-optimizer. MAP values for parameters of the full model were used to generate posterior samples of data, which were used to check the model (and its capability to recapitulate the input data), see Suppl. Fig. 2. We then proceeded with the empirical Bayes approach, keeping sample parameters fixed, and generating a full MCMC trace for each individual gene. This gave us a probability density for each of the coefficients from the design matrix, for each gene. Of this probability density, we stored 40% and 90% highest-probability density (HPD) intervals for further analyses and discarded the full trace. PROMOGEB is implemented using pymc [v5.11] (*79*) and available: (https://github.com/krooijers-sanquin/lattanzio-et-al-t-cell-activation-rbps/tree/main).

### Gene set enrichment analyses

Gene set enrichment analyses (GSEA) on modeled coefficients were performed on the MAP value after model fitting for each coefficient. We extracted gene sets from MSigDB [v2023.1.PATCH1] (*80*) and used blitzgsea [v1.3.53] (*81*) to run the analyses.

### Volcano plots and logFCs

For simple two-condition comparisons of groups of samples or replicates we employed DEP [1.24.0] (*82*). Missing values were imputed using the “manually defined left-shifted Gaussian distribution” method (shift 1.8, scale 0.3). Significance testing was performed using limma [v3.58.1] (*34*)

### Gene-editing of primary human CD8^+^ T cells

Cas9 RNP production and nucleofection was performed as described (*11*). Briefly, sgRNAs targeting RBPs (sequence in **Suppl. Table 2**) were dissolved in Nuclease Free Duplex buffer (Integrated DNA Technologies, IDT). Non-targeting negative control crRNA #1 (IDT) was mixed with tracrRNA at equimolar ratios (100uM) in nuclease-free PCR tubes and denatured at 95°C for 5 min. Nucleic acids were cooled down to RT prior to mixing and incubation for at least 10 min at RT with 30μg Alt-R™S.p.Cas9 Nuclease V3 (IDT) to generate Cas9 ribonuclear proteins (RNPs). 2×10^6^ Teff (activated for 72h) or Teff activated for 72h and then cultured for 7 days in rIL-2 containing culture medium were nucleofected with RNPs in 20μl P2 buffer (Lonza) in 16-well strips using program EH100 in a 4D Nucleofector X unit (Lonza). RBP depletion was confirmed 5d after nucleofection by immunoblot or genomic PCR, as described below.

### T cell activation

Teff cells were stimulated with 1ug/ml soluble α-CD3 (Pelicluster CD3, Sanquin) and 1 μg/mL soluble α-CD28 in culture medium for 3h. For intracellular molecule measurements, 1 μg/mL GolgiStop (BD Bioscience) and 1 μg/mL Monensin (BD Bioscience) was added during T cell activation. For translation efficiency, T cells were incubated with 5 ug/ml Puromycin dihydrochloride (Sigma) for 10 min at 37°C.

### Flow cytometry and intracellular staining

T cells were washed with FACS buffer (PBS containing 1% FBS and 2 mM EDTA) and labeled for 20 min at 4°C with α-CD4 (SK3), α-CD8 (SK1), α-CD69 (FN50), α-TIM3(73D, all BD Horizon), and α-LAG3 (11C365), α-CTLA4 (L3D10), α-CD137(4B4-1), α-PD1 (EH12.2H7), α-CD27 (O323, all Biolegend), and α-CCR7 (3D12, BD Pharmigen). CD107a (H4A3, BD Horizon) staining was performed during T cell culture for indicated time points. Dead cells were excluded with Near-IR (Life Technologies). For intracellular staining, cells were fixed and permeabilized with Cytofix/Cytoperm kit (BD Biosciences), and stained with α-IFN-γ (B27, BD Horizon), α-TNF (MAb11, BD Bioscience), α-IL2 (MQ1-17H12, BD Horizon), α-GrzB (GB11, BD Horizon), α-Puromycin (12D10, Merk), α-Perforin (RMG2a-62, Biolegend). For transcription factor staining, cells were fixed and permeabilized with eBioscience™ Foxp3/ Transcription Factor Staining Buffer Set (Invitrogen) prior to staining with α-TCF1 (C63D9, Cell Signalling), α-T-bet (4B10, BioLegend), α-FOXO1 (C29H4, Cell Signalling), α-TOX (REA473, Mintenyi), α-Blimp1 (6D3, BD), according to the manufacturer’s protocol. Data acquisition was performed using FACS Symphony A5 Cell Analyzer, data were analyzed with FlowJo (version 10.8.1, both BD Biosciences).

### Generation of *α*-CD3 expressing Mel888 target cells

A membrane-thethered anti-CD3 single-chain variable fragment (scFv) of the OKT3 clone was generated following the approach of Leitner et al. (*83*) consisting of the CD5 leader peptide, OKT3 scFv, and a leader-less CD14. Sequence is available on GenBank (ADN42857). The open-reading-frame encoding this protein and a T2A sequence followed by the red-fluorescent protein Katushka was synthesized by Twist and cloned into the pSUPER-Retro retroviral expression vector. Retroviral supernatants were generated by transient transfection of FLYRD18 cells using GeneJammer (Agilent) according to the manufacturer’s protocol. The melanoma cell line Mel888 was cultured in the presence of retrovirus-containing supernatant. After 1 week of culture, Mel888 cells were FACS-sorted based on Katushka protein expression, resulting in Mel888 cell lines with a distinct OKT3-scFv expression levels (high and low).

### T cell co-culture with tumor cells

15,000 Mel888 cells with high or low α-CD3 expression were seeded per well on a 96-well flat-bottom tissue-culture plate. After 4 hrs, 15,000 CD8^+^ T cells were added in culture medium containing 50U/ml IL-2 to achieve a 1:1 effector: target ratio. Plates were centrifuged at 10g for 1 min without brake and incubated at 37°C. For cytokine measurements: After 30 min, 1 μg/mL GolgiStop (BD Biosciences) and 1 μg/mL Monensin (BD Bioscience) was added. Cells were incubated at 37°C for 16 hrs, and cytokine production was assessed by intracellular staining (see above). For target cell killing: after 16hrs of co-culture, all adherent and non-adherent cells were collected and target cell viability was measured with Precision Count Beads (Biolegend).

### Immunoblot and Coomassie Blue Staining

Cell lysates (1×10^6^ cells/sample) were prepared by standard procedures using RIPA lysis buffer. Proteins were separated on a 4–12% SDS/PAGE and transferred onto a nitrocellulose membrane by iBlot (Thermo). Mouse monoclonal α-HuR (3A2), α-DAZAP1 (sc-373987, both Santa Cruz Biotechnology), α-PCIF1 (16082-1-AP), α-CIRBP (10209-2-AP), α-NMD3 (16060-1-AP), α-RNH1 (10345-1-AP), α-SERBP1 (10729-1-AP), α-SF3A1 (15858-1-AP), α-U2AF2 (15858-1-AP, all Proteintech) were used. As household genes, α-H3 (SC-10809, Santa Cruz), α-tubulin (T6199, Sigma), and α-RhoGDI (MAB9959, Abnova) were used. Primary antibody incubations were followed by goat a-rabbit-HRP (4050-05) or goat a-mouse-HRP (1031-05, both Southern Biotech). For silver staining, 4-12% SDS/PAGE was incubated with Coomassie Brilliant Blue staining (Merk) o/n at RT, washed with Milli-Q water to remove excess staining. Imaging was performed using ChemiDoc and Image Lab software (both Bio-Rad).

### Polymerase chain reaction (PCR)

For RBP KO PCR validation of TUT1, 100 ng genomic DNA was amplified with GoTaq G2 Flexi polymerase (Promega, M7805) with human *TUT1* specific primers, designed by using the Primer3Plus (*84*) (**Suppl. Table S3**) with the following protocol: 98°C for 5 min, 32 cycles of (30 sec at 98°C, 25 sec at 65°C, 1 min at 72°C), followed by 2 min at 72°C. PCR products were run on a 1-1.2% agarose gel. Quantification was performed with Analyze-Gel of Fiji version 2.15.1.

### RNA ligation-mediated poly(A) test (RL-PAT)

RL-PAT was performed as described (*85*). The 5’ to 5’ adenylated and 3’ blocked ‘PAT anchor’ oligo was ligated to the 3’ end of total RNA overnight at 16°C with RNA ligase 2, truncated KQ (NEB, M0373). To generate cDNA, ligated RNA was reverse transcribed with SuperScript III Reverse Transcriptase (Invitrogen, 18080044) with the ‘PAT-R1’ oligo (complementary to ‘PAT anchor’). cDNA was amplified with GoTaq G2 Flexi Polymerase (Promega, M7805), using a forward primer annealing to the 3’ UTR of the mRNA of interest (**Suppl. Table S3**) and PAT-R1 as reverse primer. All mRNA specific PAT primers were validated by performing PAT on mRNA deadenylated with oligo-d(T) and RNAse H.

### Statistical analysis and data visualization

Flow-cytometry results are shown as mean ± SD. Statistical analysis was performed in GraphPad Prism (version 9.1.1), with a two-tailed ratio paired or unpaired Student’s t-test when comparing two groups, or with Krushal-Wallis non-parametric test with Tukey-HSD correction when comparing multiple groups over different time points. p values < 0.05 were considered statistically significant. Data were visualized with GraphPad Prism (version 9.1.1). For heatmap visualization, flow cytometry data were processed in R (v4.4.1). To enable comparison across experiments, gMFI and percentage values were normalized within each donor, time point, and marker by computing log₂ (KO/control) ratios. The resulting log₂ fold-change matrix was visualized using the ComplexHeatmap package, with data scaled by row (marker) to emphasize relative differences. Hierarchical clustering was performed using Euclidean distance and complete linkage.

## Acknowledgments

We thank the lab of W. Faller and R. van der Kammen (Netherlands Cancer Institute, Amsterdam), for the use of the UV-crosslinking equipment, M. van den Biggelaar, I.P. Foskolou (Sanquin Blood Supply Foundation, Amsterdam) and the lab of R. Pillai (University of Geneva) for critical discussion, and B. Popovic (Sanquin) for critical reading of the manuscript. We thank the Wolkers lab, in particular B. Nicolet, A. Guislain (Sanquin) for technical support.

## Funding

European research council ERC-Printers 817533 (MCW)

Oncode Institute (MCW).

Landsteiner Foundation for Blood Transfusion Research (LSBR) 2202 (MCW).

## Author contributions

Conceptualization, project administration, writing: MVL, KR, MCW. Software, data analysis: KR, NS. Data curation, visualization: MVL, KR, NS, NK, NHS. Validation, investigation, methodology: MVL, AB, KB, MNM, FPJA, AJH. Funding acquisition, supervision: MCW.

## Competing interests

Authors declare no competing interests.

## Data and materials availability

TCL and OOPS proteome dataset are available at PRIDE (PXD070908)

PROMOGEB analysis pipeline is available at the following link: https://github.com/krooijers-sanquin/lattanzio-et-al-t-cell-activation-rbps/tree/main

## Supplementary Materials for

### Other Supplementary Materials for this manuscript include the following

Table S1 to S3

**Supplementary Figure 1.**
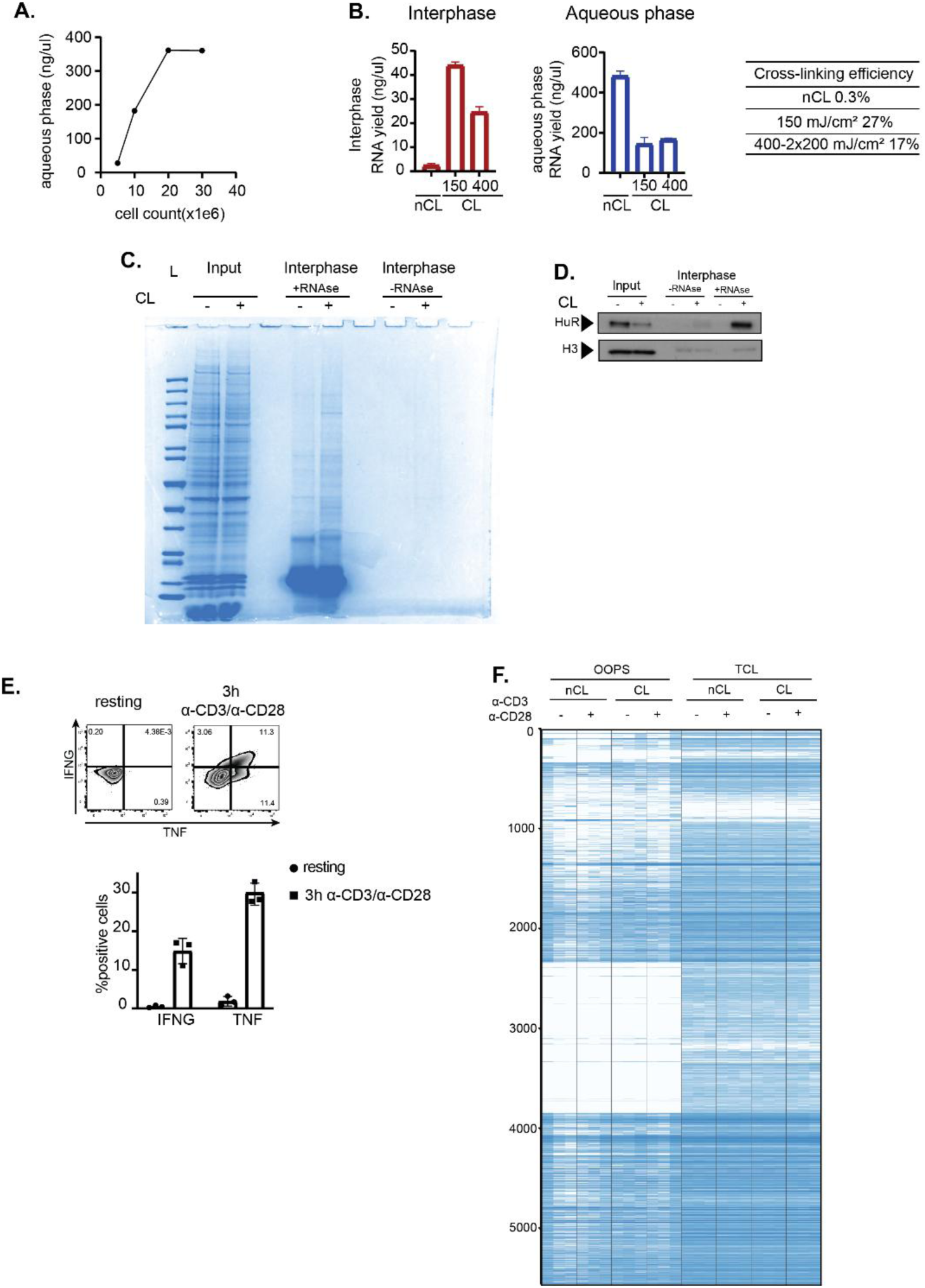
Technical validation of OOPS in human T cells. **A.** Quantification of RNA yield (ng/μl) isolated from aqueous phase per input of Teff cells. median, n=2 donors. **B.** Quantification of RNA yield (ng/μl) isolated from interphase (left) and aqueous phase (right) under different UV-cross-linking conditions (non-CL, 150mJ/mc^2^ and 400mJ/mc^2^). mean ± SD, n=2 donors. Table shows quantification of cross-linking efficiency calculated as RNA interphase (ng/μl)/ RNA (interphase + Aqueous phase) (ng/μl). **C.** Coomassie Brilliant Blue–stained gel showing protein yield from 50×10e^6^ Teff cells under different conditions: OOPS Input (nCL, CL), interphase treated with RNAse (nCL, CL) and interphase not treated with RNAse (nCL, CL). L indicates protein ladder. **D.** Immunoblot of HuR (32 kDa) and H3 (15 kDa) at indicated conditions. **E.** Representative (top) and compiled data of 3 donors (bottom) of IFN-γ and TNF production in Teff cells. mean ± SD. p≤0.05, **p≤0.01, ***p≤0.001, ****p≤0.0001 ns: non-significant. Two-tailed ratio unpaired student t-test. **F**. Raw intensity values across all samples in the proteomics dataset. Samples (in columns) are grouped by library type (OOPS, TCL), crosslinking, activation, and 3 independent replicates. Missing values are indicated in white.

**Supplementary Figure 2.**
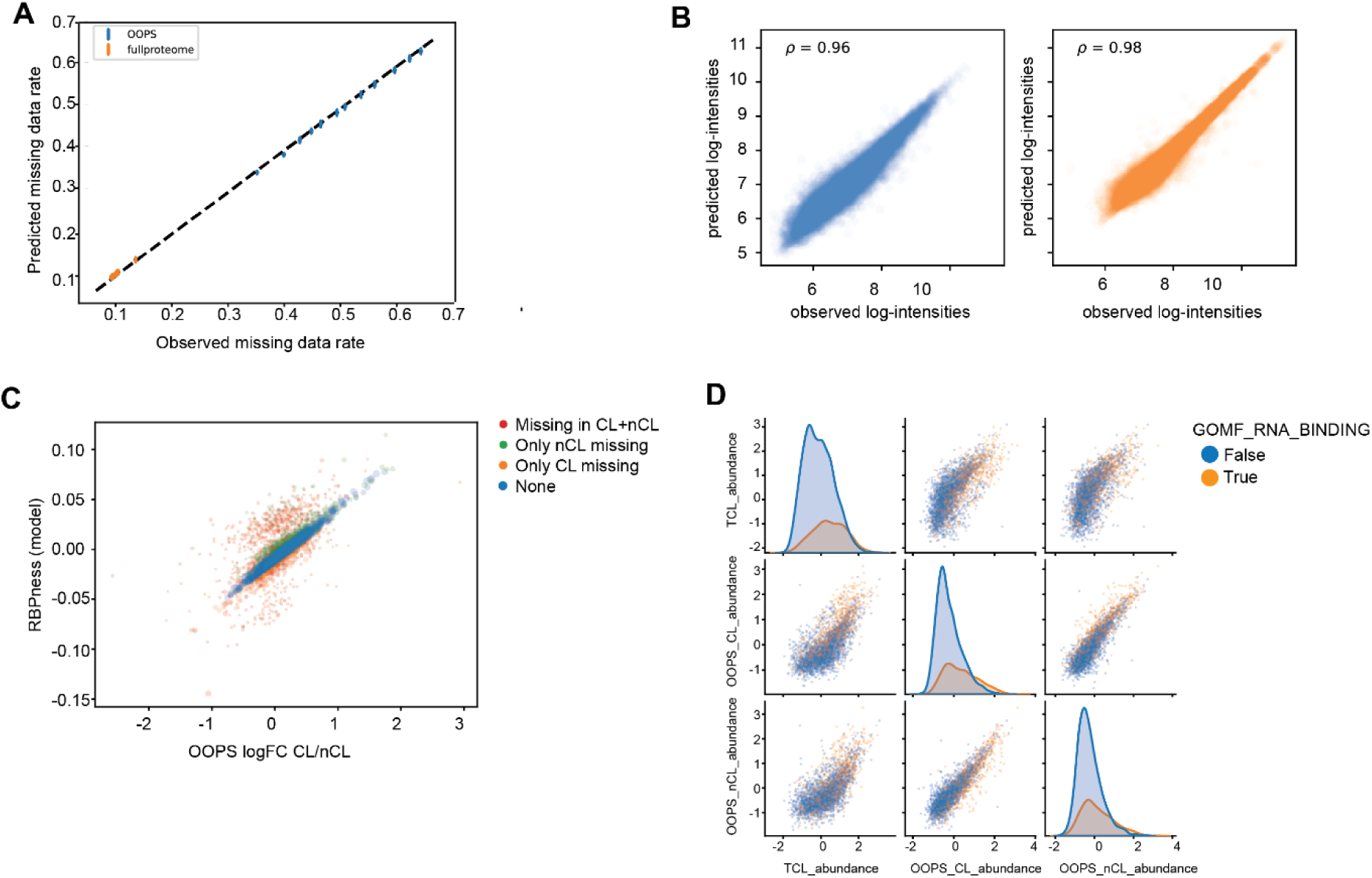
Validation of PROMOGEB models to analyse OOPS data. **A**: Predicted missing-value rates (y-axis) vs. observed missing value rates (x-axis). **B.** Predicted (y-axis) and observed (x-axis) log-intensities across samples from TCL (orange panel) or OOPS (blue panel) library types. **C**. Model coefficient for RBPness (OOPS-specific crosslinking-dependent abundance changes, y-axis) and raw log-fold-changes (x-axis). Red: proteins with missing values in both CL and nCL samples. Green: proteins with missing values in only nCL samples. Orange: proteins with missing values in only CL samples. Blue: proteins with detected abundance. **D** Correlation across Total Cell Lysate (TCL), OOPS crosslinked (CL) and OOPS non-crosslinked (nCL) samples in resting condition shows high correlation between all three sample types. Genes in orange are annotated with GO term “molecular function - RNA binding” and show elevated intensity in all three sample types (Kolmogorov-Smirnov p-value < 1e-16 in all cases).

**Supplementary Figure 3.**
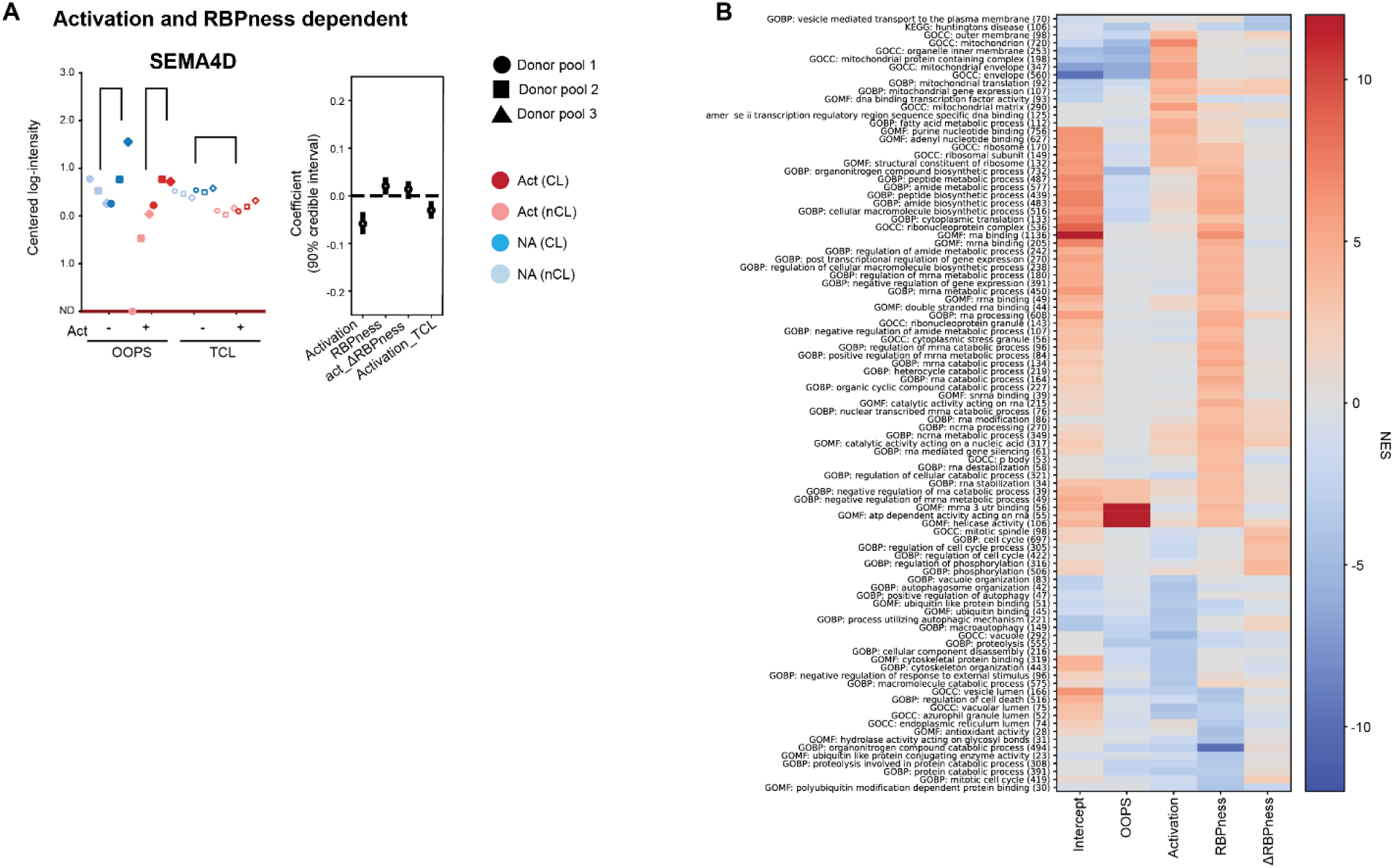
Identification of RBPs interacting with different classes of RNA. **A** SEMA4D protein abundance indicated as centered-log intensity (left panels) and biological coefficient estimates (HDIs, right panels). ND indicates “not detected”. Error bars of coefficients indicate the HDI (credible interval range) of 90%. Only biologically interpretable coefficients are shown. **B** Heatmap corresponding to Fig. 3F, showing gene sets found to be significantly enriched or depleted with p-value < 1e-3 in one of the coefficients: “OOPS specificity”, “activation”, “RBPness”, “activation-dependent RBPness (ΔRBPness)”.

**Supplementary Figure S4.**
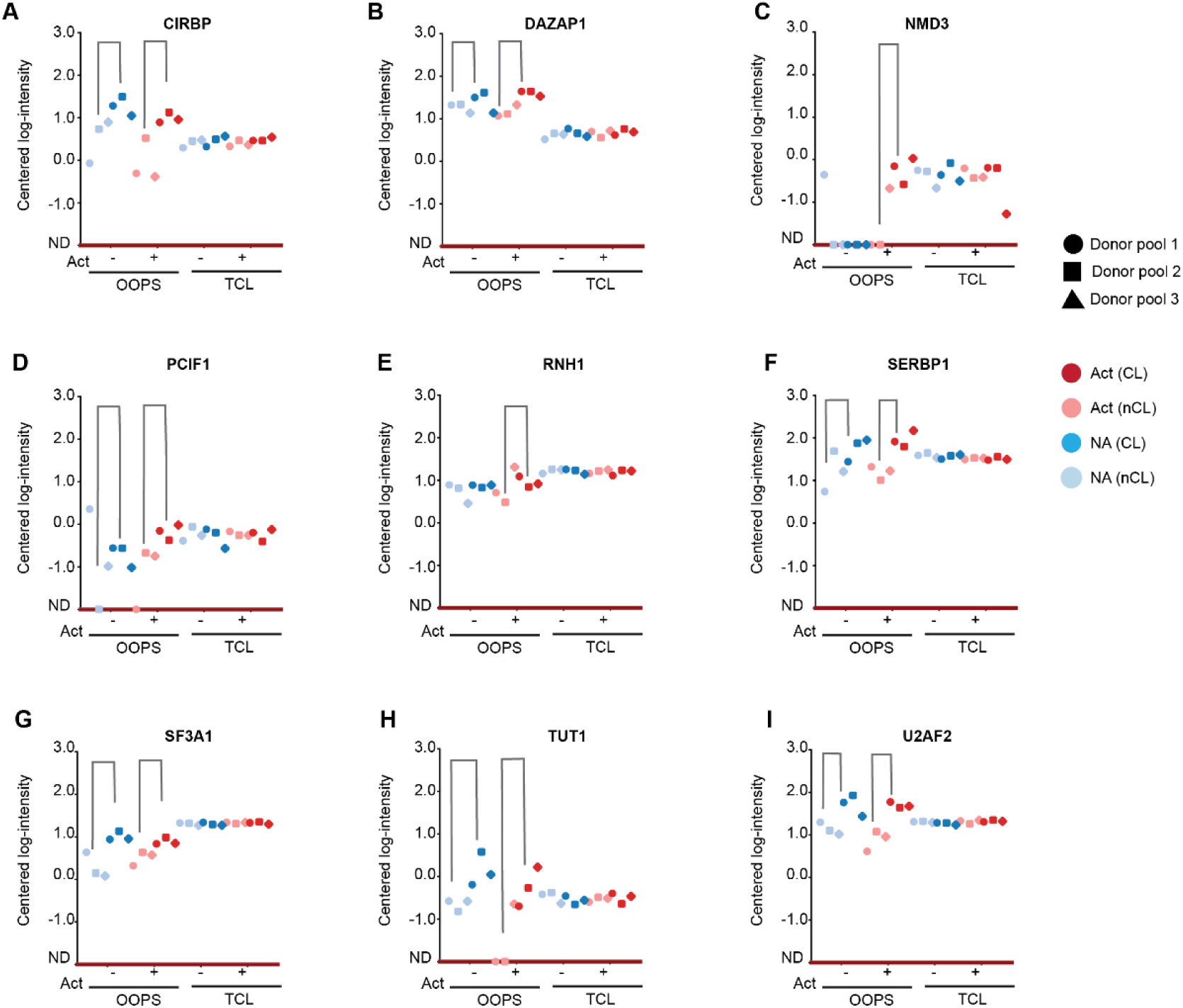
Examples of RBPs that change RBPness upon T cell activation. **A-I** Proteomics measurements of individual RBPs that were followed up by gene-editing, indicated as centered-log intensity for proteins showing changes in RNA-binding upon T cell activation.

**Supplementary Figure 5.**
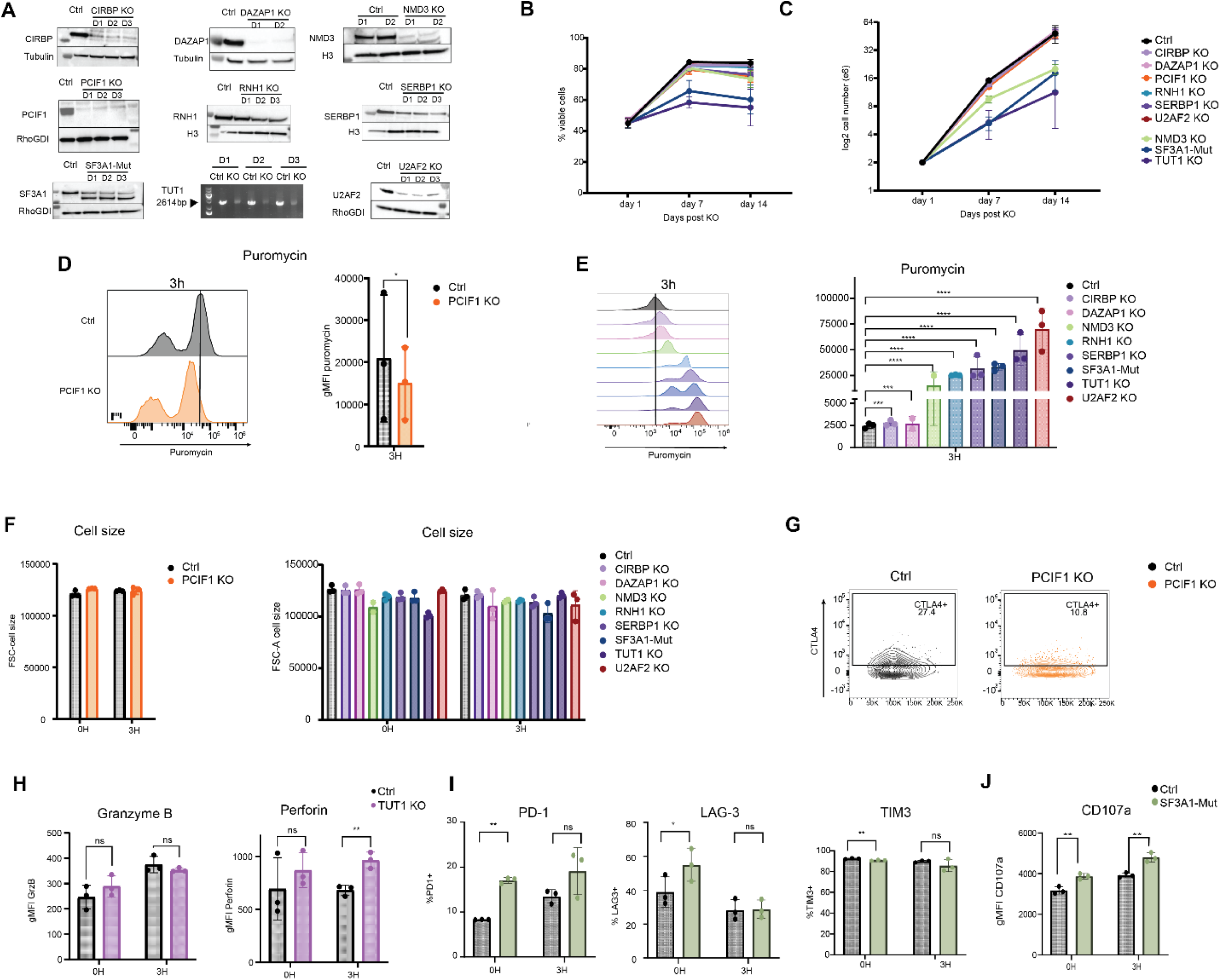
CRISPR-CAS9 gene-editing identifies RBPs modulating T cell function. **A** Validation of CRISPR-Cas9 gene editing for indicated RBPs. Efficiency was determined by immunoblot (CIRBP, DAZAP1, NMD3, PCIF1, RNH1, SERBP1, SF3A1, U2AF2), or by PCR on genomic DNA (TUT1). **B-C** Cell viability (**B**) and cell count (**C**) of indicated Teff cell type (mean ± SD, n=3; n=2 for DAZAP1 KO and NMD3 KO Teff cells). **D** Representative histogram (left) and gMFI quantification (right) of puromycin expression in 3h activated PCIF1 KO and Control Teff cells. **E** Representative histogram (left) and gMFI quantification (right) of puromycin expression in indicated Teff cells at 3h activation. **F.** Cell size quantification (FSC-A) of indicated Teff cells. **G** Representative CTLA4 expression in resting PCIF1 KO and Control Teff cells. **H** Compiled data of GMZB (left) and Perforin (right) expression in TUT1 KO and Teff cells. **I** Compiled data of PD1 (left), LAG3 (middle) and TIM3 (right) expression in SF3A1-Mut and Control Teff cells. **J** gMFI quantification of CD107a expression in SF3A1-Mut and Control Teff cells **B-J** Control, CIRBP KO, PCIF1 KO, RNH1 KO, SERBP1 KO, SF3A1-Mut, TUT1 KO, U2AF2 KO data pooled from 3 donors. DAZAP1 KO, NMD3 KO data pooled from 2 donors. Mean ± SD, p≤0.05, **p≤0.01, ***p≤0.001, ****p≤0.0001 ns: non-significant. Two-tailed ratio unpaired student t-test.

**Supplementary Figure 6.**
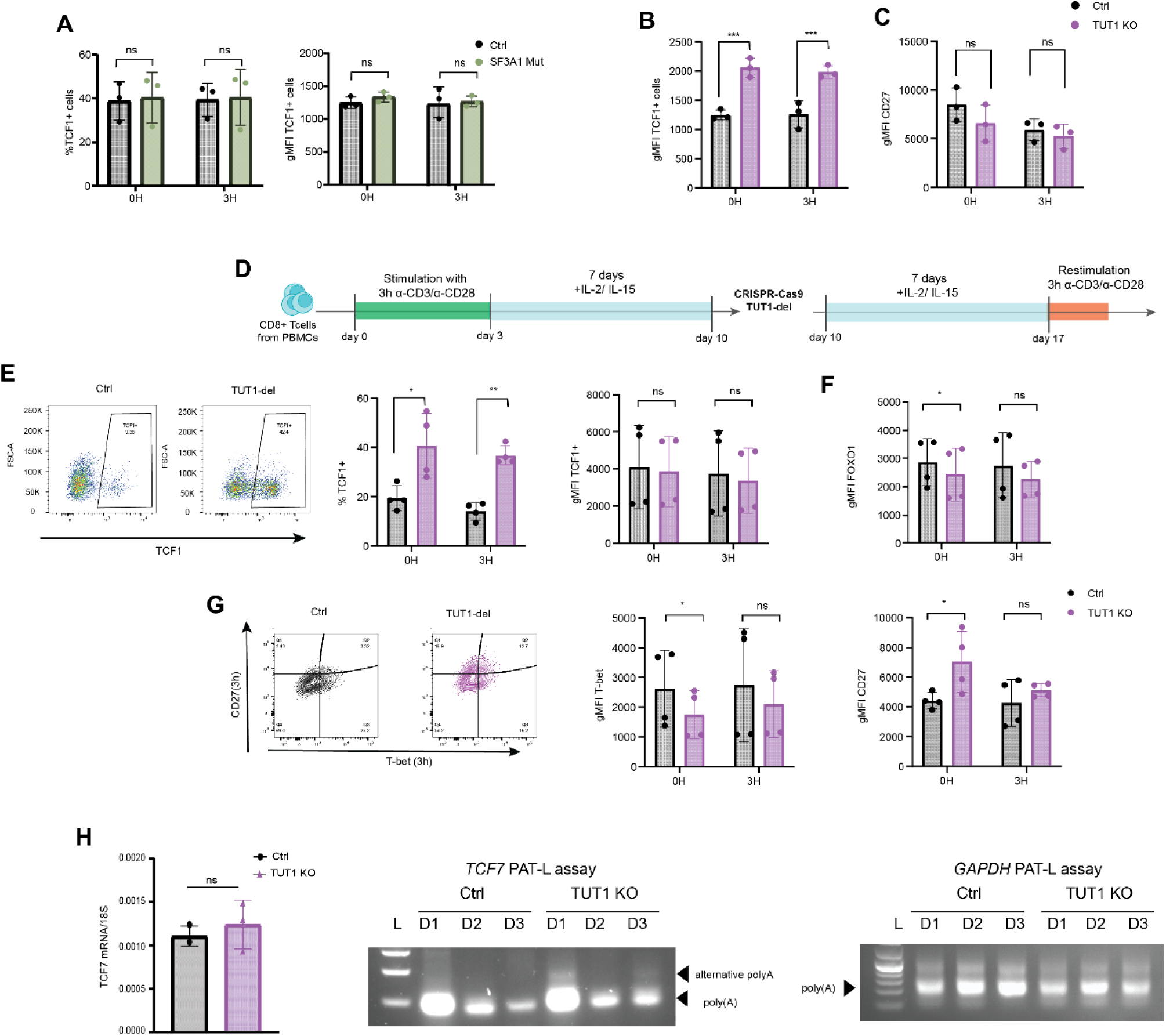
A. Effect of RBP deletion on T cell differentiation profile. **A** Compiled data of TCF1 expression in SF3A1-Mut and Control cells. **B-C** gMFI quantification of TCF1 (B) and CD27 (C) expression in TUT1 KO and Control Teff cells. **D** Schematic overview of TUT1 deletion at day 10. Teff cells generated from PBMCs by a 3-day stimulation with α-CD3/α-CD28 were cultured for 7 days before TUT1 deletion was performed. Cells were cultured for an additional 7 days, before resting and α-CD3/α-CD28 activated (3h) T cells were examined. **E-G** TCF1 expression (E), gMFI quantification of FOXO1 expression (F) and T-bet and CD27 expression in TUT1 KO and Control Teff cells. **H** Quantification (left) of *TCF7* mRNA in resting TUT1 KO and Control Teff cells. Poly(A) length measurement of *TCF7* and *GAPDH* mRNA in resting TUT1 KO and Control Teff cells, using the RL-PAT assay. Arrows indicate *TCF7* 3’UTR full-length, alternatively polyadenylated *TCF7* 3’UTR, and *GAPDH* 3’UTR full-length. **A-H** Data pooled from 3 donors. mean ± SD, p≤0.05, **p≤0.01, ***p≤0.001, ****p≤0.0001 ns: non-significant. Two-tailed ratio unpaired student t-test.

**Supplementary Figure 7.**
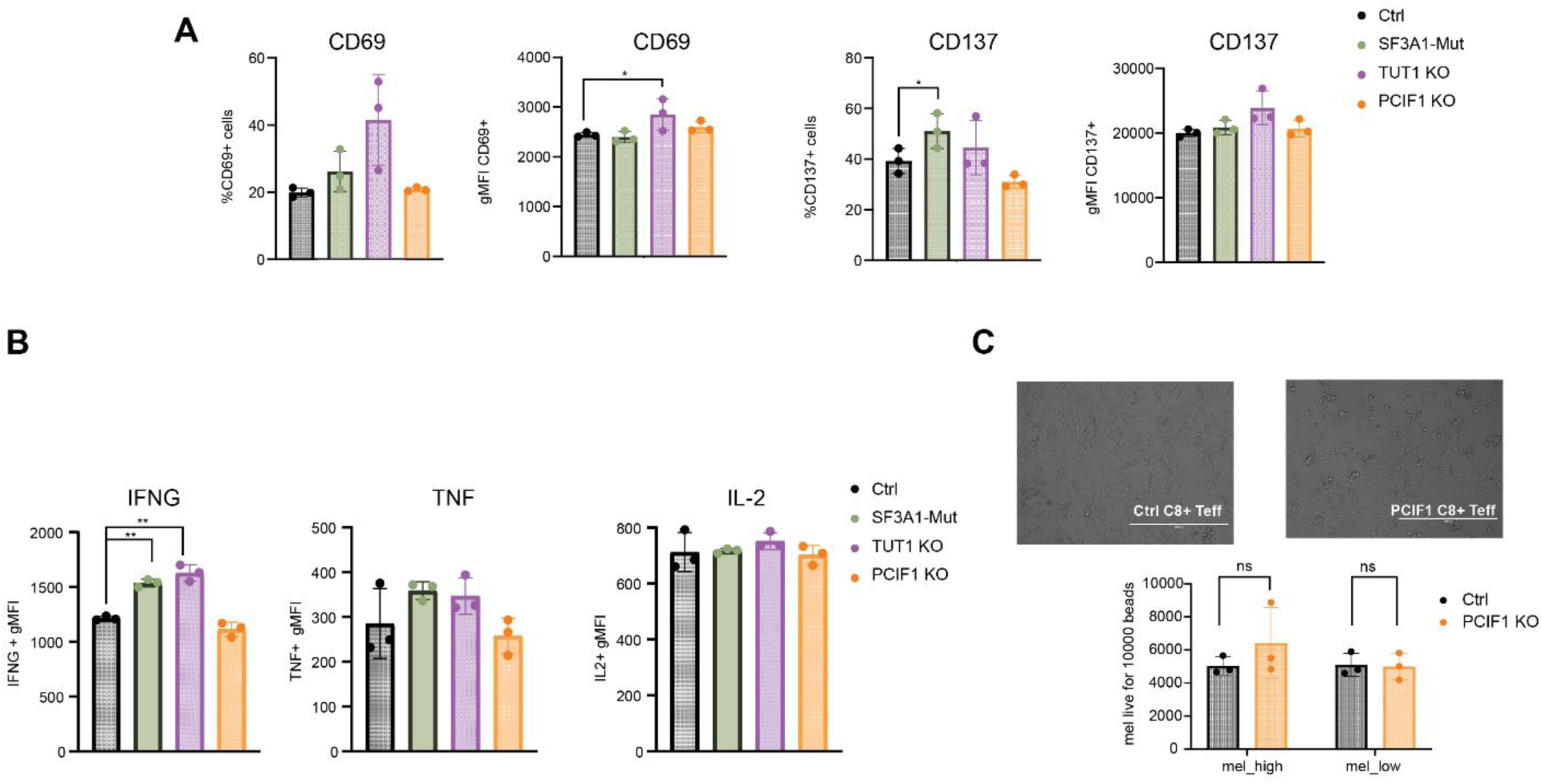
RBP KO T cell phenotype in co-culture system. **A** Compiled data of CD69 (left) and CD137 (right) expression in indicated Teff cells co-cultured with α-CD3 high Mel888 for 16h. **B** gMFI quantification of IFN-γ (left), TNF (middle) and IL2 (right) in indicated Teff cells co-cultured with α-CD3 medium Mel888 for 16h in the presence of GolgiStop and Monensin added after 1h. **C** Representative image (top) of 16hrs co-culture of Control and PCIF1 KO Teff cells with α-CD3 high Mel888 cells (scale bar = 400μm). Quantification (bottom) of live Mel888 cells per 10.000 counting beads. **A-C** Data pooled from 3 donors. mean ± SD, p≤0.05, **p≤0.01, ***p≤0.001, ****p≤0.0001 ns: non-significant. Two-tailed ratio unpaired student t-test.

## Notes

https://github.com/krooijers-sanquin/2024-proteomics-preprocessing

### Competing Interest Statement

The authors have declared no competing interest.

## References

1. T. Wolf, W. Jin, G. Zoppi, I. A. Vogel, M. Akhmedov, C. K. E. Bleck, T. Beltraminelli, J. C. Rieckmann, N. J. Ramirez, M. Benevento, S. Notarbartolo, D. Bumann, F. Meissner, B. Grimbacher, M. Mann, A. Lanzavecchia, F. Sallusto, I. Kwee, R. Geiger, Dynamics in protein translation sustaining T cell preparedness. Nat Immunol 21, 927–937 (2020).

2. M. V. Lattanzio, N. Šoštarić, N. Kanagasabesan, B. Popović, A. Bradarić, L. Wardak, A. Guislain, P. Savakis, E. Tutucci, M. C. Wolkers, Single-molecule imaging of transcription dynamics, RNA localization and fate in human T cells. EMBO J, 1–18 (2025).

3. F. Salerno, S. Engels, M. van den Biggelaar, F. P. J. van Alphen, A. Guislain, W. Zhao, D. L. Hodge, S. E. Bell, J. P. Medema, M. von Lindern, M. Turner, H. A. Young, M. C. Wolkers, Translational repression of pre-formed cytokine-encoding mRNA prevents chronic activation of memory T cells. Nature Immunology 2018 19:8 19, 828–837 (2018).

4. B. P. Nicolet, M. C. Wolkers, The relationship of mRNA with protein expression in CD8+ T cells associates with gene class and gene characteristics. PLoS One 17 (2022).

5. H. Weerakoon, A. Mohamed, Y. Wong, J. Chen, B. Senadheera, O. Haigh, T. S. Watkins, S. Kazakoff, P. Mukhopadhyay, J. Mulvenna, J. J. Miles, M. M. Hill, A. Lepletier, Integrative temporal multi-omics reveals uncoupling of transcriptome and proteome during human T cell activation. npj Systems Biology and Applications 2024 10:1 10, 1–13 (2024).

6. F. Salerno, M. Turner, M. C. Wolkers, Dynamic Post-Transcriptional Events Governing CD8+ T Cell Homeostasis and Effector Function. Trends in immunology 41.3 240–245 (2020).

7. D. L. Kontoyiannis, An RNA checkpoint that keeps immunological memory at bay. Nature Immunology 2018 19:8 19, 795–797 (2018).

8. M. Turner, M. D. DÍaz-Muñoz, RNA-binding proteins control gene expression and cell fate in the immune system. Nat Immunol 19, 120–129 (2018).

9. M. W. Hentze, P. Sommerkamp, V. Ravi, F. Gebauer, Rethinking RNA-binding proteins: Riboregulation challenges prevailing views. Cell 188, 4811–4827 (2025).

10. G. Petkau, T. J. Mitchell, K. Chakraborty, S. E. Bell, V. D’Angeli, L. Matheson, D. J. Turner, A. Saveliev, O. Gizlenci, F. Salerno, P. D. Katsikis, M. Turner, The timing of differentiation and potency of CD8 effector function is set by RNA binding proteins. Nature Communications 2022 13:1 13, 1–16 (2022).

11. B. Popović, B. P. Nicolet, A. Guislain, S. Engels, A. P. Jurgens, N. Paravinja, J. J. Freen-van Heeren, F. P. J. van Alphen, M. van den Biggelaar, F. Salerno, M. C. Wolkers, Time-dependent regulation of cytokine production by RNA binding proteins defines T cell effector function. Cell Rep 42, 112419 (2023).

12. K. P. Hoefig, A. Reim, C. Gallus, E. H. Wong, G. Behrens, C. Conrad, M. Xu, L. Kifinger, T. Ito-Kureha, K. A. Y. Defourny, A. Geerlof, J. Mautner, S. M. Hauck, D. Baumjohann, R. Feederle, M. Mann, M. Wierer, E. Glasmacher, V. Heissmeyer, Defining the RBPome of primary T helper cells to elucidate higher-order Roquin-mediated mRNA regulation. Nature Communications 2021 12:1 12, 1–18 (2021).

13. T. Nakahama, Y. Kato, J. I. Kim, T. Vongpipatana, Y. Suzuki, C. R. Walkley, Y. Kawahara, ADAR 1-mediated RNA editing is required for thymic self-tolerance and inhibition of autoimmunity. EMBO Rep 19 (2018).

14. M. E. Cook, T. R. Bradstreet, A. M. Webber, J. Kim, A. Santeford, K. M. Harris, M. K. Murphy, J. Tran, N. M. Abdalla, E. A. Schwarzkopf, S. C. Greco, C. M. Halabi, R. S. Apte, P. J. Blackshear, B. T. Edelson, The ZFP36 family of RNA binding proteins regulates homeostatic and autoreactive T cell responses. Sci Immunol 7 (2022).

15. M. J. Moore, N. E. Blachere, J. J. Fak, C. Y. Park, K. Sawicka, S. Parveen, I. Zucker-Scharff, B. Moltedo, A. Y. Rudensky, R. B. Darnell, ZFP36 RNA-binding proteins restrain T cell activation and anti-viral immunity. Elife 7 (2018).

16. Y. Yao, Y. Yang, W. Guo, L. Xu, M. You, Y. C. Zhang, Z. Sun, X. Cui, G. Yu, Z. Qi, J. Liu, F. Wang, J. Liu, T. Zhao, L. Ye, Y. G. Yang, S. Yu, METTL3-dependent m6A modification programs T follicular helper cell differentiation. Nature Communications 2021 12:1 12, 1–16 (2021).

17. H. B. Li, J. Tong, S. Zhu, P. J. Batista, E. E. Duffy, J. Zhao, W. Bailis, G. Cao, L. Kroehling, Y. Chen, G. Wang, J. P. Broughton, Y. G. Chen, Y. Kluger, M. D. Simon, H. Y. Chang, Z. Yin, R. A. Flavell, m6A mRNA methylation controls T cell homeostasis by targeting the IL-7/STAT5/SOCS pathways. Nature 2017 548:7667 548, 338–342 (2017).

18. D. Mai, O. Johnson, J. Reff, T. J. Fan, J. Scholler, N. C. Sheppard, C. H. June, Combined disruption of T cell inflammatory regulators Regnase-1 and Roquin-1 enhances antitumor activity of engineered human T cells. Proc Natl Acad Sci U S A 120, e2218632120 (2023).

19. T. Mino, Y. Murakawa, A. Fukao, A. Vandenbon, H. H. Wessels, D. Ori, T. Uehata, S. Tartey, S. Akira, Y. Suzuki, C. G. Vinuesa, U. Ohler, D. M. Standley, M. Landthaler, T. Fujiwara, O. Takeuchi, Regnase-1 and roquin regulate a common element in inflammatory mRNAs by spatiotemporally distinct mechanisms. Cell 161, 1058–1073 (2015).

20. F. Gebauer, T. Schwarzl, J. Valcárcel, M. W. Hentze, RNA-binding proteins in human genetic disease. Nature Reviews Genetics 2020 22:3 22, 185–198 (2020).

21. H. Schmidt, T. Raj, T. J. O’Neill, A. Muschaweckh, F. Giesert, A. Negraschus, K. P. Hoefig, G. Behrens, L. Esser, C. Baumann, R. Feederle, C. Plaza-Sirvent, A. Geerlof, A. Gewies, S. E. Isay, J. Ruland, I. Schmitz, W. Wurstd, T. Korn, D. Krappmann, V. Heissmeyer, Unrestrained cleavage of Roquin-1 by MALT1 induces spontaneous T cell activation and the development of autoimmunity. Proc Natl Acad Sci U S A 120, e2309205120 (2023).

22. K. Matsushita, O. Takeuchi, D. M. Standley, Y. Kumagai, T. Kawagoe, T. Miyake, T. Satoh, H. Kato, T. Tsujimura, H. Nakamura, S. Akira, Zc3h12a is an RNase essential for controlling immune responses by regulating mRNA decay. Nature 2009 458:7242 458, 1185–1190 (2009).

23. S. J. Tavernier, V. Athanasopoulos, P. Verloo, G. Behrens, J. Staal, D. J. Bogaert, L. Naesens, M. De Bruyne, S. Van Gassen, E. Parthoens, J. Ellyard, J. Cappello, L. X. Morris, H. Van Gorp, G. Van Isterdael, Y. Saeys, M. Lamkanfi, P. Schelstraete, J. Dehoorne, V. Bordon, R. Van Coster, B. N. Lambrecht, B. Menten, R. Beyaert, C. G. Vinuesa, V. Heissmeyer, M. Dullaers, F. Haerynck, A human immune dysregulation syndrome characterized by severe hyperinflammation with a homozygous nonsense Roquin-1 mutation. Nat Commun 10 (2019).

24. Y. Li, S. P. Vyas, I. Mehta, N. Asada, I. Dey, T. C. Taylor, R. Bechara, N. Amatya, F. E. Y. Aggor, B. M. Coleman, D. D. Li, K. Yamamoto, O. Ezenwa, Y. Sun, E. Sterneck, C. J. McManus, U. Panzer, P. S. Biswas, R. Savan, J. Das, S. L. Gaffen, The RNA binding protein Arid5a drives IL-17–dependent autoantibody-induced glomerulonephritis. Journal of Experimental Medicine 221 (2024).

25. G. Assoni, V. La Pietra, R. Digilio, C. Ciani, N. V. Licata, M. Micaelli, E. Facen, W. Tomaszewska, L. Cerofolini, A. Pérez-Ràfols, M. Varela Rey, M. Fragai, A. Woodhoo, L. Marinelli, D. Arosio, I. Bonomo, A. Provenzani, P. Seneci, HuR-targeted agents: An insight into medicinal chemistry, biophysical, computational studies and pharmacological effects on cancer models. Adv Drug Deliv Rev 181 (2022).

26. Y. Hua, T. A. Vickers, H. L. Okunola, C. F. Bennett, A. R. Krainer, Antisense Masking of an hnRNP A1/A2 Intronic Splicing Silencer Corrects SMN2 Splicing in Transgenic Mice. Am J Hum Genet 82, 834–848 (2008).

27. Y. Hua, K. Sahashi, F. Rigo, G. Hung, G. Horev, C. F. Bennett, A. R. Krainer, Peripheral SMN restoration is essential for long-term rescue of a severe spinal muscular atrophy mouse model. Nature 478, 123–126 (2011).

28. J. Wei, L. Long, W. Zheng, Y. Dhungana, S. A. Lim, C. Guy, Y. Wang, Y. D. Wang, C. Qian, B. Xu, A. Kc, J. Saravia, H. Huang, J. Yu, J. G. Doench, T. L. Geiger, H. Chi, Targeting Regnase-1 programs long-lived effector T cells for cancer therapy. Nature 576, 471 (2019).

29. N. D. Zandhuis, A. Guislain, A. Popalzij, S. Engels, B. Popović, M. Turner, M. C. Wolkers, Regulation of IFN-γ production by ZFP36L2 in T cells is time-dependent. Eur J Immunol 54, 2451018 (2024).

30. A. P. Jurgens, J. Zwijnen, F. P. J. van Alphen, A. Bradarić, K. Rooijers, A. J. Hoogendijk, B. Popović, M. C. Wolkers, mTOR signaling promotes cytokine production in T cells through 3’UTR-mediated translation control. bioRxiv, 2024.10.01.616022 (2024).

31. E. L. Van Nostrand, P. Freese, G. A. Pratt, X. Wang, X. Wei, R. Xiao, S. M. Blue, J. Y. Chen, N. A. L. Cody, D. Dominguez, S. Olson, B. Sundararaman, L. Zhan, C. Bazile, L. P. B. Bouvrette, J. Bergalet, M. O. Duff, K. E. Garcia, C. Gelboin-Burkhart, M. Hochman, N. J. Lambert, H. Li, M. P. McGurk, T. B. Nguyen, T. Palden, I. Rabano, S. Sathe, R. Stanton, A. Su, R. Wang, B. A. Yee, B. Zhou, A. L. Louie, S. Aigner, X. D. Fu, E. Lécuyer, C. B. Burge, B. R. Graveley, G. W. Yeo, A large-scale binding and functional map of human RNA-binding proteins. Nature 2020 583:7818 583, 711–719 (2020).

32. R. M. L. Queiroz, T. Smith, E. Villanueva, M. Marti-Solano, M. Monti, M. Pizzinga, D. M. Mirea, M. Ramakrishna, R. F. Harvey, V. Dezi, G. H. Thomas, A. E. Willis, K. S. Lilley, Comprehensive identification of RNA–protein interactions in any organism using orthogonal organic phase separation (OOPS). Nat Biotechnol 37, 169–178 (2019).

33. J. I. Perez-Perri, B. Rogell, T. Schwarzl, F. Stein, Y. Zhou, M. Rettel, A. Brosig, M. W. Hentze, Discovery of RNA-binding proteins and characterization of their dynamic responses by enhanced RNA interactome capture. Nature Communications 2018 9:1 9, 1–13 (2018).

34. M. E. Ritchie, B. Phipson, D. Wu, Y. Hu, C. W. Law, W. Shi, G. K. Smyth, limma powers differential expression analyses for RNA-sequencing and microarray studies. Nucleic Acids Res 43, e47–e47 (2015).

35. M. Li, G. K. Smyth, Neither random nor censored: estimating intensity-dependent probabilities for missing values in label-free proteomics. Bioinformatics 39 (2023).

36. E. C. Urdaneta, C. H. Vieira-Vieira, T. Hick, H. H. Wessels, D. Figini, R. Moschall, J. Medenbach, U. Ohler, S. Granneman, M. Selbach, B. M. Beckmann, Purification of cross-linked RNA-protein complexes by phenol-toluol extraction. Nature Communications 2019 10:1 10, 1–17 (2019).

37. M. Caudron-Herger, S. F. Rusin, M. E. Adamo, J. Seiler, V. K. Schmid, E. Barreau, A. N. Kettenbach, S. Diederichs, R-DeeP: Proteome-wide and Quantitative Identification of RNA-Dependent Proteins by Density Gradient Ultracentrifugation. Mol Cell 75, 184–199.e10 (2019).

38. J. Trendel, T. Schwarzl, R. Horos, A. Prakash, A. Bateman, M. W. Hentze, J. Krijgsveld, The Human RNA-Binding Proteome and Its Dynamics during Translational Arrest. Cell 176, 391–403.e19 (2019).

39. F. Salerno, N. A. Paolini, R. Stark, M. Von Lindern, M. C. Wolkers, Distinct PKC-mediated posttranscriptional events set cytokine production kinetics in CD8+ T cells. Proc Natl Acad Sci U S A 114, 9677–9682 (2017).

40. J. C. Schwartz, C. C. Ebmeier, E. R. Podell, J. Heimiller, D. J. Taatjes, T. R. Cech, FUS binds the CTD of RNA polymerase II and regulates its phosphorylation at Ser2. Genes Dev 26, 2690–2695 (2012).

41. E. Izaurralde, J. Lewis, C. Gamberi, A. Jarmolowski, C. McGuigan, I. W. Mattaj, A cap-binding protein complex mediating U snRNA export. Nature 376, 709–712 (1995).

42. Y. Takagaki, J. L. Manley, RNA recognition by the human polyadenylation factor CstF. Mol Cell Biol 17, 3907–3914 (1997).

43. P. C. Lin, R. M. Xu, Structure and assembly of the SF3a splicing factor complex of U2 snRNP. EMBO Journal 31, 1579–1590 (2012).

44. R. Choudhury, S. G. Roy, Y. S. Tsai, A. Tripathy, L. M. Graves, Z. Wang, The splicing activator DAZAP1 integrates splicing control into MEK/Erk-regulated cell proliferation and migration. Nat Commun 5, 3078 (2014).

45. D. L. Mellman, M. L. Gonzales, C. Song, C. A. Barlow, P. Wang, C. Kendziorski, R. A. Anderson, A PtdIns4,5P2-regulated nuclear poly(A) polymerase controls expression of select mRNAs. Nature 2008 451:7181 451, 1013–1017 (2008).

46. Y. Takagaki, J. L. Manley, RNA recognition by the human polyadenylation factor CstF. Mol Cell Biol 17, 3907–3914 (1997).

47. A. P. Sudheesh, N. Mohan, N. Francis, R. S. Laishram, R. A. Anderson, Star-PAP controlled alternative polyadenylation coupled poly(A) tail length regulates protein expression in hypertrophic heart. Nucleic Acids Res 47, 10771–10787 (2019).

48. R. S. Laishram, R. A. Anderson, The poly A polymerase Star-PAP controls 3′-end cleavage by promoting CPSF interaction and specificity toward the pre-mRNA. EMBO J 29, 4132 (2010).

49. T. Kimura, I. Hashimoto, T. Nagase, J. I. Fujisawa, CRM1-dependent, but not ARE-mediated, nuclear export of IFN-α1 mRNA. J Cell Sci 117, 2259–2270 (2004).

50. T. C. Walther, M. Fornerod, H. Pickersgill, M. Goldberg, T. D. Allen, I. W. Mattaj, The nucleoporin Nup153 is required for nuclear pore basket formation, nuclear pore complex anchoring and import of a subset of nuclear proteins. EMBO J 20, 5703–5714 (2001).

51. E. Sendinc, D. Valle-Garcia, A. Dhall, H. Chen, T. Henriques, J. Navarrete-Perea, W. Sheng, S. P. Gygi, K. Adelman, Y. Shi, PCIF1 Catalyzes m6Am mRNA Methylation to Regulate Gene Expression. Mol Cell 75, 620–630.e9 (2019).

52. S. Akichika, S. Hirano, Y. Shichino, T. Suzuki, H. Nishimasu, R. Ishitani, A. Sugita, Y. Hirose, S. Iwasaki, O. Nureki, T. Suzuki, Cap-specific terminal N 6 -methylation of RNA by an RNA polymerase II–associated methyltransferase. Science (1979) 363 (2019).

53. J. Ayache, M. Bénard, M. Ernoult-Lange, N. Minshall, N. Standart, M. Kress, D. Weil, P-body assembly requires DDX6 repression complexes rather than decay or Ataxin2/2L complexes. Mol Biol Cell 26, 2579–2595 (2015).

54. K. Araki, M. Morita, A. G. Bederman, B. T. Konieczny, H. T. Kissick, N. Sonenberg, R. Ahmed, Translation is actively regulated during the differentiation of CD8 + effector T cells. Nat Immunol 18, 1046–1057 (2017).

55. F. Dragon, P. A. Compagnone-Post, B. M. Mitchell, K. A. Porwancher, K. A. Wehner, S. Wormsley, R. E. Settlage, J. Shabanowitz, Y. Osheim, A. L. Beyer, D. F. Hunt, S. J. Baserga, A large nucleolar U3 ribonucleoprotein required for 18S ribosomal RNA biogenesis. Nature 417, 967–970 (2002).

56. L. Lee, J. B. Whittall, WDR75: An essential protein for ribosome assembly undergoing purifying selection. PLoS One 20, e0318395 (2025).

57. J. H.-N. Ho, A. W. Johnson, NMD3 Encodes an Essential Cytoplasmic Protein Required for Stable 60S Ribosomal Subunits in Saccharomyces cerevisiae. Mol Cell Biol 19, 2389–2399 (1999).

58. A. G. Malyutin, S. Musalgaonkar, S. Patchett, J. Frank, A. W. Johnson, Nmd3 is a structural mimic of eIF 5A, and activates the cp GTP ase Lsg1 during 60S ribosome biogenesis. EMBO J 36, 854–868 (2017).

59. R. Allam, V. Chennupati, D. F. T. Veiga, K. M. Maslowski, A. Tardivel, M. Quadroni, M. Duchosal, H. R. MacDonald, N. Fasel, A. Angelillo-Scherrer, P. Schneider, T. Hoang, An Unexpected Role for Ribonuclease Inhibitor (RNH1) in Erythropoiesis. Blood 124, 244–244 (2014).

60. M. Stillinovic, M. A. Sarangdhar, N. Andina, A. Tardivel, F. Greub, G. Bombaci, C. Ansermet, M. Zatti, D. Saha, J. Xiong, T. Sagae, M. Yokogawa, M. Osawa, M. Heller, A. Keogh, I. Keller, A. Angelillo-Scherrer, R. Allam, Ribonuclease inhibitor and angiogenin system regulates cell type–specific global translation. Sci Adv 10 (2024).

61. A. P. Jurgens, B. Popović, M. C. Wolkers, T cells at work: How post-transcriptional mechanisms control T cell homeostasis and activation. European Journal of Immunology 51.9 2178–2187 (2021).

62. N. Nameki, M. Takizawa, T. Suzuki, S. Tani, N. Kobayashi, T. Sakamoto, Y. Muto, K. Kuwasako, Structural basis for the interaction between the first SURP domain of the SF3A1 subunit in U2 snRNP and the human splicing factor SF1. Protein Science 31, e4437 (2022).

63. B. Xiang, M. Zhang, K. Li, Z. Zhang, Y. Liu, M. Gao, X. Wang, X. Xiao, Y. Sun, C. He, J. Shi, H. Fan, X. Xing, G. Xu, Y. Yao, G. Chen, H. Zhu, C. Yi, J. Zhang, The epitranscriptional factor PCIF1 orchestrates CD8+ T cell ferroptosis and activation to control antitumor immunity. Nat Immunol 26, 252–264 (2025).

64. R. R. Pandey, E. Delfino, D. Homolka, A. Roithova, K. M. Chen, L. Li, G. Franco, C. B. Vågbø, E. Taillebourg, M. O. Fauvarque, R. S. Pillai, The Mammalian Cap-Specific m6Am RNA Methyltransferase PCIF1 Regulates Transcript Levels in Mouse Tissues. Cell Rep 32 (2020).

65. A. M. Intlekofer, N. Takemoto, E. J. Wherry, S. A. Longworth, J. T. Northrup, V. R. Palanivel, A. C. Mullen, C. R. Gasink, S. M. Kaech, J. D. Miller, L. Gapin, K. Ryan, A. P. Russ, T. Lindsten, J. S. Orange, A. W. Goldrath, R. Ahmed, S. L. Reiner, Effector and memory CD8+ T cell fate coupled by T-bet and eomesodermin. Nat Immunol 6, 1236–1244 (2005).

66. K. J. Oestreich, A. C. Huang, A. S. Weinmann, The lineage-defining factors T-bet and Bcl-6 collaborate to regulate Th1 gene expression patterns. Journal of Experimental Medicine 208, 1001–1013 (2011).

67. J. Hendriks, L. A. Gravestein, K. Tesselaar, R. A. W. Van Lier, T. N. M. Schumacher, J. Borst, CD27 is required for generation and long-term maintenance of T cell immunity. Nat Immunol 1, 433–440 (2000).

68. R. H. Michelini, A. L. Doedens, A. W. Goldrath, S. M. Hedrick, Differentiation of CD8 memory T cells depends on Foxo1. J Exp Med 210, 1189–1200 (2013).

69. K. Bresser, B. Popović, M. C. Wolkers, What’s in a name: the multifaceted function of DNA- and RNA-binding proteins in T cell responses. FEBS Journal 292, 1853–1867 (2025).

70. W. H. Hudson, E. A. Ortlund, The structure, function and evolution of proteins that bind DNA and RNA. Nat Rev Mol Cell Biol 15, 749–760 (2014).

71. L. Merendino, S. Guth, D. Bilbao, C. Martínez, J. Valcárcel, Inhibition of msl-2 splicing by Sex-lethal reveals interaction between U2AF35 and the 3’ splice site AG. Nature 402, 838–841 (1999).

72. A. Crisci, F. Raleff, I. Bagdiul, M. Raabe, H. Urlaub, J. C. Rain, A. Krämer, Mammalian splicing factor SF1 interacts with SURP domains of U2 snRNP-associated proteins. Nucleic Acids Res 43, 10456 (2015).

73. D. Blake, C. M. Radens, M. B. Ferretti, M. R. Gazzara, K. W. Lynch, Alternative splicing of apoptosis genes promotes human T cell survival. Elife 11 (2022).

74. B. Yang Yang, J.-F. Chang, J. R. Parnes, C. Garrison Fathman, “T Cell Receptor (TCR) Engagement Leads to Activation-induced Splicing of Tumor Necrosis Factor (TNF) Nuclear Pre-mRNA” The Journal of experimental medicine 188.2 247–254 (1998).

75. J. A. Best, D. A. Blair, J. Knell, E. Yang, V. Mayya, A. Doedens, M. L. Dustin, A. W. Goldrath, P. Monach, S. A. Shinton, R. R. Hardy, R. Jianu, D. Koller, J. Collins, R. Gazit, B. S. Garrison, D. J. Rossi, K. Narayan, K. Sylvia, J. Kang, A. Fletcher, K. Elpek, A. Bellemare-Pelletier, D. Malhotra, S. Turley, J. A. Best, V. Jojic, D. Koller, T. Shay, A. Regev, N. Cohen, P. Brennan, M. Brenner, T. Kreslavsky, N. A. Bezman, J. C. Sun, C. C. Kim, L. L. Lanier, J. Miller, B. Brown, M. Merad, E. L. Gautier, C. Jakubzick, G. J. Randolph, F. Kim, T. N. Rao, A. Wagers, T. Heng, M. Painter, J. Ericson, S. Davis, A. Ergun, M. Mingueneau, D. Mathis, C. Benoist, Transcriptional insights into the CD8 + T cell response to infection and memory T cell formation. Nat Immunol (2013).

76. J. T. Chang, E. J. Wherry, A. W. Goldrath, Molecular regulation of effector and memory T cell differentiation. Nature immunology 15.12 1104–1115 (2014).

77. K. Boulias, D. Toczydłowska-Socha, B. R. Hawley, N. Liberman, K. Takashima, S. Zaccara, T. Guez, J. J. Vasseur, F. Debart, L. Aravind, S. R. Jaffrey, E. L. Greer, Identification of the m6Am Methyltransferase PCIF1 Reveals the Location and Functions of m6Am in the Transcriptome. Mol Cell 75, 631–643.e8 (2019).

78. J. Mauer, X. Luo, A. Blanjoie, X. Jiao, A. V. Grozhik, D. P. Patil, B. Linder, B. F. Pickering, J. J. Vasseur, Q. Chen, S. S. Gross, O. Elemento, F. Debart, M. Kiledjian, S. R. Jaffrey, Reversible methylation of m6Am in the 5′ cap controls mRNA stability. Nature 2016 541:7637 541, 371–375 (2016).

79. O. Abril-Pla, V. Andreani, C. Carroll, L. Dong, C. J. Fonnesbeck, M. Kochurov, R. Kumar, J. Lao, C. C. Luhmann, O. A. Martin, M. Osthege, R. Vieira, T. Wiecki, R. Zinkov, PyMC: a modern, and comprehensive probabilistic programming framework in Python. PeerJ Comput Sci 9, e1516 (2023).

80. A. Liberzon, A. Subramanian, R. Pinchback, H. Thorvaldsdóttir, P. Tamayo, J. P. Mesirov, Molecular signatures database (MSigDB) 3.0. Bioinformatics 27, 1739–1740 (2011).

81. A. Lachmann, Z. Xie, A. Ma’ayan, blitzGSEA: efficient computation of gene set enrichment analysis through gamma distribution approximation. Bioinformatics 38, 2356–2357 (2022).

82. X. Zhang, A. H. Smits, G. B. A. Van Tilburg, H. Ovaa, W. Huber, M. Vermeulen, Proteome-wide identification of ubiquitin interactions using UbIA-MS. Nat Protoc 13, 530–550 (2018).

83. J. Leitner, W. Kuschei, K. Grabmeier-Pfistershammer, R. Woitek, E. Kriehuber, O. Majdic, G. Zlabinger, W. F. Pickl, P. Steinberger, T cell stimulator cells, an efficient and versatile cellular system to assess the role of costimulatory ligands in the activation of human T cells. J Immunol Methods 362, 131–141 (2010).

84. S. Rozen, H. Skaletsky, Primer3 on the WWW for General Users and for Biologist Programmers. Methods Mol Biol 132, 365–386 (2000).

85. H. A. Meijer, M. Bushell, K. Hill, T. W. Gant, A. E. Willis, P. Jones, C. H. de Moor, A novel method for poly(A) fractionation reveals a large population of mRNAs with a short poly(A) tail in mammalian cells. Nucleic Acids Res 35, e132 (2007).

